# PHALCON: phylogeny-aware variant calling from large-scale single-cell panel sequencing datasets

**DOI:** 10.1101/2024.12.26.630385

**Authors:** Priya, Sunkara B V Chowdary, Aditya Gautam, Hamim Zafar

## Abstract

Single-cell sequencing (SCS) enables variant detection and tumor phylogeny reconstruction for resolving intra-tumor heterogeneity (ITH), which causes drug resistance and cancer relapse. Recently emerged panel sequencing methods sequence disease-specific genes across thousands of cells, but existing variant callers and SCS-specific phylogenetic methods struggle with large-scale datasets and amplification biases in panel-based sequencing protocols. We present a statistical variant caller, PHALCON for scalable mutation detection from large-scale single-cell panel sequencing data by modeling tumor evolution under a finite-sites model along a clonal phylogeny. Across a wide variety of simulation settings, PHALCON outperformed state-of-the-art methods in variant calling, tumor phylogeny inference, and runtime. From triple negative breast cancer (TNBC) and acute myeloid leukemia (AML) datasets, PHALCON detected novel somatic mutations with high functional impact, resolved clonal substructure and rare clones. In AML, PHALCON also uncovered poor-survival subgroups harboring DNA methylation and chromatin/cohesin mutations and revealed novel cellular co-occurrence and exclusivity patterns of driver mutations.

## Introduction

Since the inception of the single-cell sequencing (SCS) technologies [1], diverse fields of biology including developmental biology, neurobiology, immunology, and cancer biology have greatly benefited from outstanding advancements in the fields of single-cell genomics and transcriptomics [2, 3, 4, 5, 6]. The genomic cellular profiles generated by such technologies allow for investigating genetic variability across cells at an unprecedented resolution, and thus can unravel the somatic evolutionary events in normal as well as pathological development. Single-cell genomic profiles are particularly beneficial for elucidating the intra-tumor heterogeneity (ITH), which can play an important role in developing therapeutic resistance and relapse [7], in addition to driving cancer progression and metastasis [8]. ITH is caused by the presence of multiple genetically distinct cellular populations (clones), which arise due to the ongoing somatic evolution and selection process in the cancer tissue [9]. Accurate detection of the mutations of the clones as well as the inference of the chronological order of the mutations is crucial for understanding ITH which can further aid in developing precision medicine. Even though SCS has the power and resolution to resolve ITH [6], the aforementioned tasks become difficult due to elevated error rates associated with the whole-genome amplification (WGA) process required for SCS library preparation. WGA technologies while necessary for amplifying the small quantity of DNA material available in an individual cell to provide sufficient DNA for sequencing, they result in several technical artifacts, including allelic dropout (ADO), false-positive (FP) and false-negative (FN) errors, and coverage non-uniformity [10].

As traditional variant callers [11] were ill-suited for handling SCS-specific technical errors, new variant callers such as Monovar [12] and SCcaller [13] were developed for detecting point mutations from SCS datasets. In another direction, a suite of tumor phylogeny inference methods [14, 15, 16, 17] were developed for the reconstruction of evolutionary history of the cells from the noisy single nucleotide variant (SNV) profiles obtained from SCS datasets. Variant calling and tumor phylogeny inference were combined in methods such as SCIPhi [18] and scVILP [19], which showed that the joint inference of cellular phylogeny and SNVs further improves variant calling from SCS datasets. Phylovar [20] extended the likelihood-based approach for phylogeny-aware detection of SNVs for single-cell whole genome sequencing (scWGS) datasets consisting of large number of genomic loci.

However, all these phylogeny-aware variant callers worked under the infinite sites assumption (ISA), which simplifies modeling by assuming that a given mutation cannot recur in the phylogeny and that mutations, once acquired, are not lost. ISA was found to be violated in real tumor samples [21] and several single-cell tumor phylogeny inference methods [15, 16] showed the superiority of finite-sites model of evolution in capturing different mutational events. Following this, more recent phylogeny-aware SCS variant callers such as SCIPhIN [22] and SIEVE [23] also allowed for the loss and recurrence of mutations on the phylogeny in their underlying variant calling formulation. One of the major drawbacks of existing single-cell variant callers is that they can handle low-throughput datasets consisting of only hundreds of cells. Recently, high-throughput microfluidic-based SCS platforms (e.g., Tapestri by Mission Bio) have enabled the mutation profiling of thousands of cells in parallel where a panel of known disease-specific genes [24, 25, 26] or a custom gene panel [27] is targeted. Existing variant calling methods are not capable of handling thousands of cells present in such datasets and they often fail to generate any result. Moreover, the existing methods are optimized for small-scale SCS datasets and they can produce a large number of false positives when applied on the panel sequencing datasets for which the underlying error rates may be different from the small-scale datasets. Finally, the usage of cell-resolution phylogeny in the discovery and genotyping of variants as done by some state-of-the-art methods is intractable for datasets with thousands of cells.

To address these challenges, we present PHALCON, a novel statistical phylogeny-aware variant calling method that enables scalable mutation detection from large-scale single-cell panel sequencing data consisting of thousands of cells by modeling their evolutionary history under a finite-sites model along a clonal phylogeny. For each cell and each site in the panel sequencing dataset, PHALCON computes genotype likelihoods using beta-binomial distributions whose parameters are learned using a maximum likelihood approach. Based on the genotype likelihoods of candidate variant sites, PHALCON infers the clonal clusters using graph-based clustering and reconstructs a clonal phylogeny and the most likely mutation history using a likelihood-based framework that maximizes the likelihood of the observed read counts given the genotypes. Using a large number of simulated datasets representing a wide range of experimental settings, we showed that PHALCON outperforms existing state-of-the-art methods in terms of variant calling accuracy, accuracy in inferring the tumor phylogeny, and runtime. Finally, we applied PHALCON to real tumor single-cell st cancer (TNBC) patients [27] and a large cohort (123 patients) of acute myeloid leukemia (AML) patients [24], where PHALCON detected novel somatic mutations in important oncogenes, tumor suppressor genes, and AML driver genes respectively, with high functional impact and orthogonal support in bulk datasets. PHALCON was able to provide a more comprehensive understanding of clonal substructure and evolution in TNBC by resolving a larger number of cellular subpopulations, identifying rare clones, and elucidating lineages harboring deleterious mutations in key oncogenes and tumor suppressor genes. For AML, PHALCON identified subgroups of patients with poor survival harboring DNA methylation and chromatin/cohesin mutations and also unraveled novel cellular-level co-occurrence and mutual exclusivity patterns of driver gene mutations in major pathways associated with AML.

## Results

### Overview of PHALCON

We have developed PHALCON, a phylogeny-aware likelihood-based framework for performing scalable variant calling on high-throughput single-cell panel sequencing datasets consisting of thousands of cells (Fig. 1a). PHALCON employs a finite-sites evolution model that accounts for heterozygous mutation as well as recurrent mutations (parallel mutation and mutation loss) on the branches of a tumor phylogeny (Fig. 1b). PHALCON infers somatic SNVs and small indels (Fig. 1c) and genotypes them in individual cells by reconstructing a clonal phylogeny of the cells where the clonal substructure of the cells and the finite-sites model of evolution together guide the variant calling and genotyping process.

**Figure 1:**
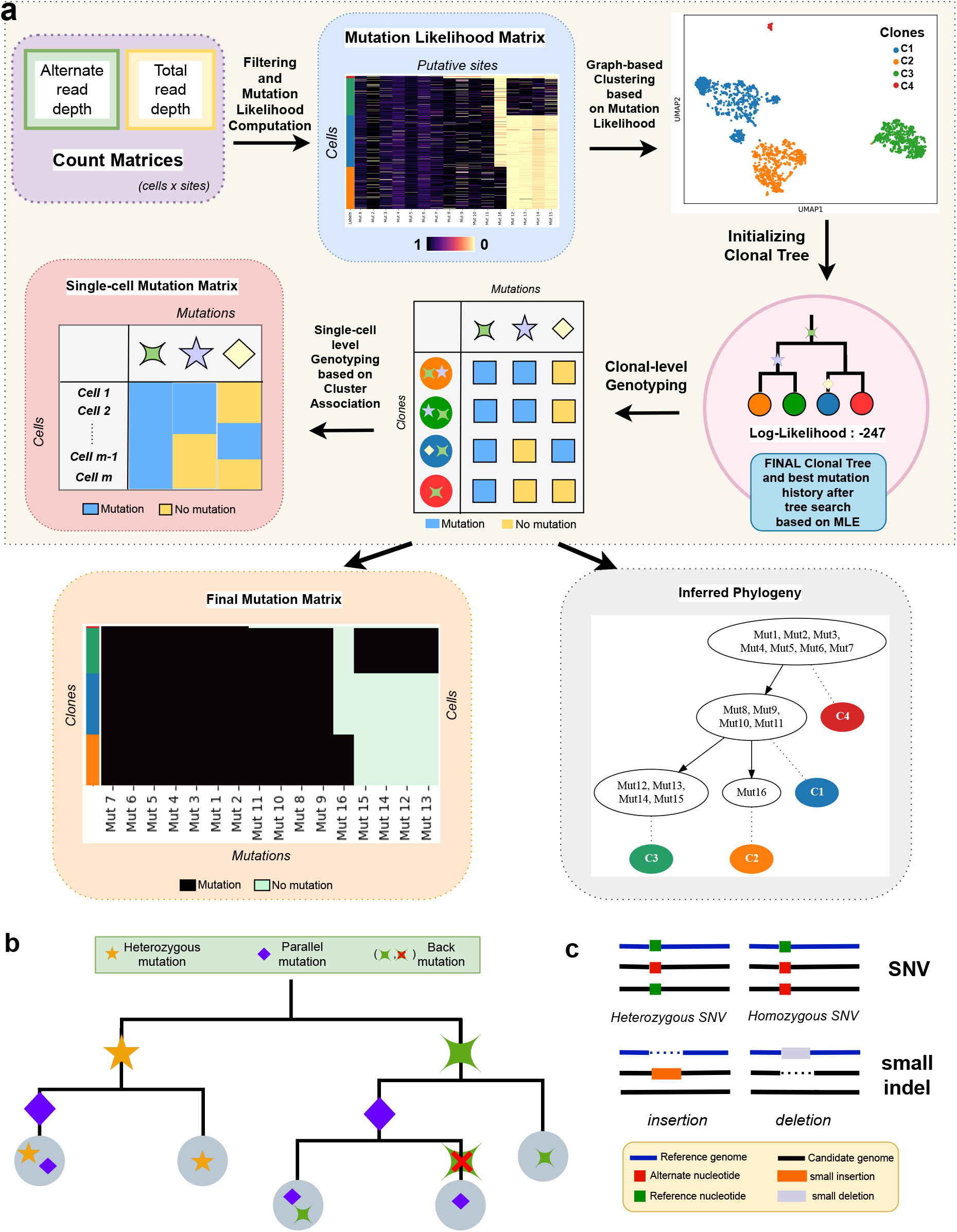
Overview of PHALCON. (a) PHALCON is a phylogeny-aware statistical model that performs variant calling from large-scale single-cell sequencing datasets using a likelihood-based framework. PHALCON first identifies candidate mutation sites by removing uninformative sites using a set of filters. It then calculates a cells × sites mutation likelihood matrix, using which PHALCON performs graph-based clustering and clusters the cells into their respective clones. PHALCON then performs a likelihood-based search to infer the optimal underlying clonal phylogeny as well as the clonal genotypes. PHALCON finally genotypes the mutated sites in single cells using the clonal genotypes and clonal membership of cells. Final output of PHALCON consists of the cell-level genotype matrix delineating the presence or absence of mutations in each cell as well as the clonal membership of each cell and a clonal phylogeny whose branches are annotated with the mutations. (b) Types of mutational events modeled by PHALCON’s finite-sites model of evolution. (c) PHALCON can infer single nucleotide variants (SNVs) and small indels from single-cell read count dataset.

PHALCON uses as input the alternate and total read counts for each site in each cell using which it computes likelihood of mutation at each site and each cell using beta-binomial distributions whose parameters are learned using a maximum likelihood approach. After that it identifies the putative variant sites using a multi-step filtering algorithm that removes information from low-quality reads, uninformative reads, and potential false positives. Using the mutation likelihood matrix for the putative variant sites, PHALCON applies graph-based clustering (spectral clustering by default) to infer *K* cellular sub-populations in the form of clonal clusters of cells. With the inferred clonal clusters as leaves, PHALCON initializes a clonal phylogeny and performs a stochastic search in the tree search space for inferring a clonal phylogeny and the most likely mutation history using a likelihood-based framework that maximizes the log-likelihood of the observed read counts given the genotypes (see Methods for details). While inferring the best possible mutation history on a given clonal phylogeny, PHALCON considers all possible mutation histories associated with the finite-sites evolution model and maximizes a penalized log-likelihood function (see Methods) which uses a regulariser on the number of occurrences of recurrent mutations to avoid overfitting. Once PHALCON infers the optimal clonal phylogeny and the best mutation history of all sites, we genotype the clonal clusters. Consequently, the cells are genotyped as per their clonal membership.

### Evaluation on simulated data

We first evaluated the variant calling performance of PHALCON using a comprehensive set of simulated datasets encompassing a wide variety of experimental settings and compared it against that of the most recent state-of-the-art SCS variant callers, namely SCIPhIN [22] and Phylovar [20]. Furthermore, we tested another state-of-the-art variant caller, SIEVE [23] which failed to produce any result for any of our datasets in spite of an exhaustive runtime of 72 hours and, hence, was not included in our benchmarking. We simulated datasets by expanding the simulator introduced in [18] to resemble data generated by panel sequencing platforms (e.g., Tapestri by Mission Bio) (see Methods). The datasets were simulated across six different scenarios: i) varying number of cells, ii) varying number of clones, iii) varying rates of mutation loss and recurrence, iv) varying ADO rate, v) varying FP rate, and vi) varying rates of copy number alteration. The accuracy of variant calling was evaluated based on F1 score which measures the harmonic mean of precision and recall (see Methods).

We first investigated how the methods perform as the number of cells increases in the dataset by simulating datasets with 2000, 5000, and 10000 cells. For these datasets, 50 mutation sites were distributed across 10 clones. Both SCIPhIN and Phylovar failed to yield any result within a 72-hour time frame for 5000 and 10000 cells datasets making PHALCON the only method capable of performing variant calling for the large-scale panel sequencing datasets (Fig. 2a). For datasets with 2000 cells, for which SCIPhIN and Phylovar were able to produce results, PHALCON demonstrated superior performance in terms of F1 score compared to both the methods (47.5% and 8.15% improvement over SCIPhIN and Phylovar respectively). PHALCON’s improved F1 score was resulted by improved recall as well as much improved precision as both Phylovar and SCIPhIN inferred a large number of false positives (∼ 60 times more putative sites were inferred as compared to the actual number of mutation sites, Supplementary Fig. 1). Since SCIPhIN and Phylovar could only produce results for 2000 cells, for the rest of the experiments the number of cells was set to 2000, number of mutation sites was set to 50 and other parameters were varied. For varying number of clones, *k* ∈ {7, 10, 15}, PHALCON outperformed the other methods in all settings by achieving 36.33-47.54% and 7.47-8.15% improvement in F1 score over SCIPhIN and Phylovar respectively (Fig. 2a). Next, we wanted to evaluate the ability of the methods in handling increased rates of mutation losses and recurrence. We generated datasets under three conditions - increased rate of back mutation, increased rate of parallel mutation, and increased rate of both back and parallel mutations (see Supplementary Table 1). For each scenario, PHALCON achieved higher precision, recall, and F1 score as compared to SCIPhIN and Phylovar (Fig. 2a). For the experiments involving elevated error rates, the performance of SCIPhIN and Phylovar degraded with increase in ADO or FP rate. In comparison, PHALCON was robust to increase in error rates and had a consistent performance across increased error rates (Fig. 2a, 47.19-51.67% and 7.5-19.01% improvement in F1 score as compared to SCIPhIN and Phylovar). Since PHALCON does not model copy number events which can have significant effect on the read count of mutated loci, we next investigated its performance in variant calling and genotyping in the presence of additional wild type alleles. PHALCON demonstrated a fairly consistent performance across varying levels of copy number events (25% and 50% mutated loci harboring copy number alterations) and outperformed SCIPhIN and Phylovar in all settings (Fig. 2a). With increasing copy number events, while PHALCON showed a stable performance, performance of SCIPhIN and Phylovar dropped more quickly. We further tested PHALCON’s ability to handle data devoid of mutation losses and recurrence by simulating datasets under the infinite sites model of evolution for which PHALCON achieved higher precision, recall, and F1 score as compared to SCIPhIN and Phylovar (Supplementary Fig. 2a-c).

**Figure 2:**
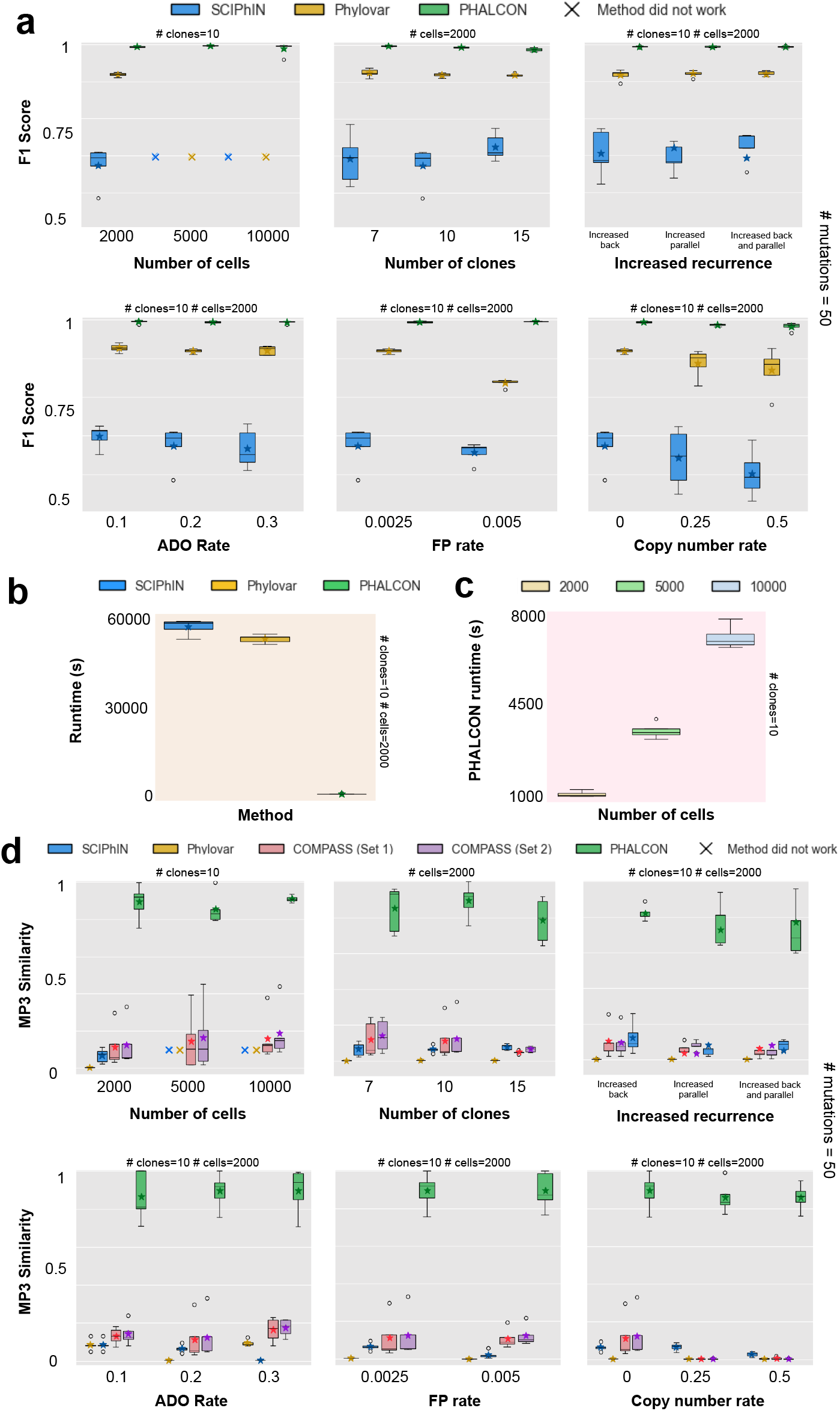
Benchmarking of PHALCON on simulated datasets. (a) Comparison of F1 score between SCIPhIN, Phylovar and PHALCON across various experimental settings for evaluating accuracy of mutation calling. (b) Runtime comparison between SCIPhIN, Phylovar and PHALCON for datasets consisting of 2000 cells. (c) Runtime of PHALCON with increasing number of cells.(d) Comparison of MP3 similarity between SCIPhIN, Phylovar, COMPASS (Set 1 and Set 2) and PHALCON across various experimental settings to evaluate the accuracy of the inferred phylogeny. In (a) and (d), experimental settings include varying number of cells, varying number of clones, varying rates of mutation loss and recurrence, varying ADO and FP rates, and copy number alteration rate. The simulated datasets contained 50 mutations across all experiments. In these figures, the boxplots depict the median and the interquartile range (IQR, the range between the 25^*th*^ and 75^*th*^ percentile); stars indicate means; whiskers extend to 1.5 times the interquartile range; and outliers beyond this range are denoted as empty circles. A ‘×’ denotes instances where the method failed to produce any result within 72 hours.

Apart from low accuracy, large runtime is a significant challenge faced by the existing variant callers when encountered with large-scale panel sequencing datasets. PHALCON exhibited a massive improvement (44 − 47.5×) in runtime over SCIPhIN and Phylovar. For the datasets with 2000 cells, PHALCON had an average runtime of ∼ 22.3 minutes as compared to 17.43 hours and 16.2 hours for that of SCIPhIN and Phylovar respectively (Fig. 2b). PHALCON’s runtime also increased linearly with the number of cells where it had average runtime of 64 minutes and 125.7 minutes for 5000 and 10000 cells respectively (Fig. 2c).

We further assessed the accuracy of PHALCON in reconstructing the evolutionary history using multipoly-occurring labels triplet-based (MP3) similarity [28], a metric that quantifies phylogenetic similarity, even among phylogenies with disparate sets of mutations. For phylogeny inference, apart from SCIPhIN and Phylovar, we also compared PHALCON against the state-of-the-art phylogeny inference method, COMPASS [29], which can infer a joint phylogeny of SNVs and CNAs from targeted large-scale SCS datasets. Since COMPASS requires the read count information of the variant sites as input, we passed the read count information from the ground truth mutation sites as input to COMPASS. However, the phylogeny inferred by COMPASS only contained a subset of the mutation sites passed as input. To account for this, we computed MP3 score for COMPASS under two settings - considering all the ground truth mutation sites (Set 1) and considering only the mutation sites retained by COMPASS in the phylogenetic tree (Set 2). Across all simulated datasets across all experimental settings, PHALCON outperformed all other methods in reconstructing the phylogenetic tree by large margins (575.85-25623.8%, 841.06-32931.78%, and 410.43-31463.3% improvement in MP3 similarity over SCIPhIN, Phylovar, and COMPASS respectively) (Fig. 2d). Even for datasets generated under the infinite sites model or harboring copy number events, PHALCON majorly outperformed the other methods (Supplementary Fig. 2d).

### PHALCON detects novel somatic mutations with functional impact in triple-negative breast cancer

We applied PHALCON to a cohort of 5 untreated triple-negative breast cancer (TNBC) patients [27] for which multi-patient-targeted (MPT) scDNA-seq was performed on a total of 23526 cells for 330 targeted sites selected based on bulk deep-exome sequencing. In the original study, the authors used the Mission Bio scDNA-seq platform to generate the datasets. PHALCON detected a total of 226 somatic mutations across all samples (on average ∼ 45 mutations/sample). Two samples (TN4 and TN5) had higher mutation burden (*>* 50 mutations) as compared to the other three samples (TN1, TN2 and TN3). Figure 3a shows the top 10 mutated genes with non-silent mutations across the five samples. Out of these 10 genes, 7 were reported to be mutated in the original study, including tumor suppressor genes *TP53, RASAL2*, and *NOTCH3*. PHALCON identified *TP53* to be mutated in all the samples and all these mutations occurred in the DNA binding domain (Fig. 3b) of *TP53*, which is a known hotspot of oncogenic somatic mutations [30]. In *RASAL2*, which is known to promote TNBC progression [31], PHALCON identified a novel deleterious mutation (not reported in the original study) in the catalytic RasGAP domain (Fig. 3b). In the original study, *NOTCH3* was reported to be mutated in only one sample, in contrast, PHALCON detected *NOTCH3* mutation in three samples, two of which were clonal and one was sub-clonal (Fig. 3a). PHALCON further inferred mutations in genes that were not reported to be mutated in the original study. These included mutations in *PROCR*, a cell surface protein known to be enriched in TNBCs [32], *CNOT3*, a tumor suppressor gene and *PRIM2*. Interestingly, *PROCR* was found to harbor a clonal missense mutation in all five samples, and the mutation was predicted to have a high functional impact by CADD (Supplementary Fig. 3a). Pathogenicity evaluation using MutPred2 [33] resulted in a high pathogenicity score for the *PROCR* mutation with multiple molecular mechanisms altered (Supplementary Fig. 3b). To evaluate the importance of *PROCR* in TNBC, we analyzed single-cell RNA-seq data from epithelial cells of 10 TNBC patients, and found the expression of *PROCR* to be higher in cancer epithelial cells as compared to normal epithelial cells (Supplementary Figs. 3c-d). Survival analysis with the TCGA cohort (89 TNBC patients) [34] showed poor survival for the patients harboring higher expression of *PROCR*, indicating its importance in TNBC progression (Supplementary Fig. 3e). For the novel missense mutation in *CNOT3* detected by PHALCON, MutPred2 predicted alterations in six molecular mechanisms resulting in high pathogenicity (score = 0.737) (Supplementary Fig. 3f).

**Figure 3:**
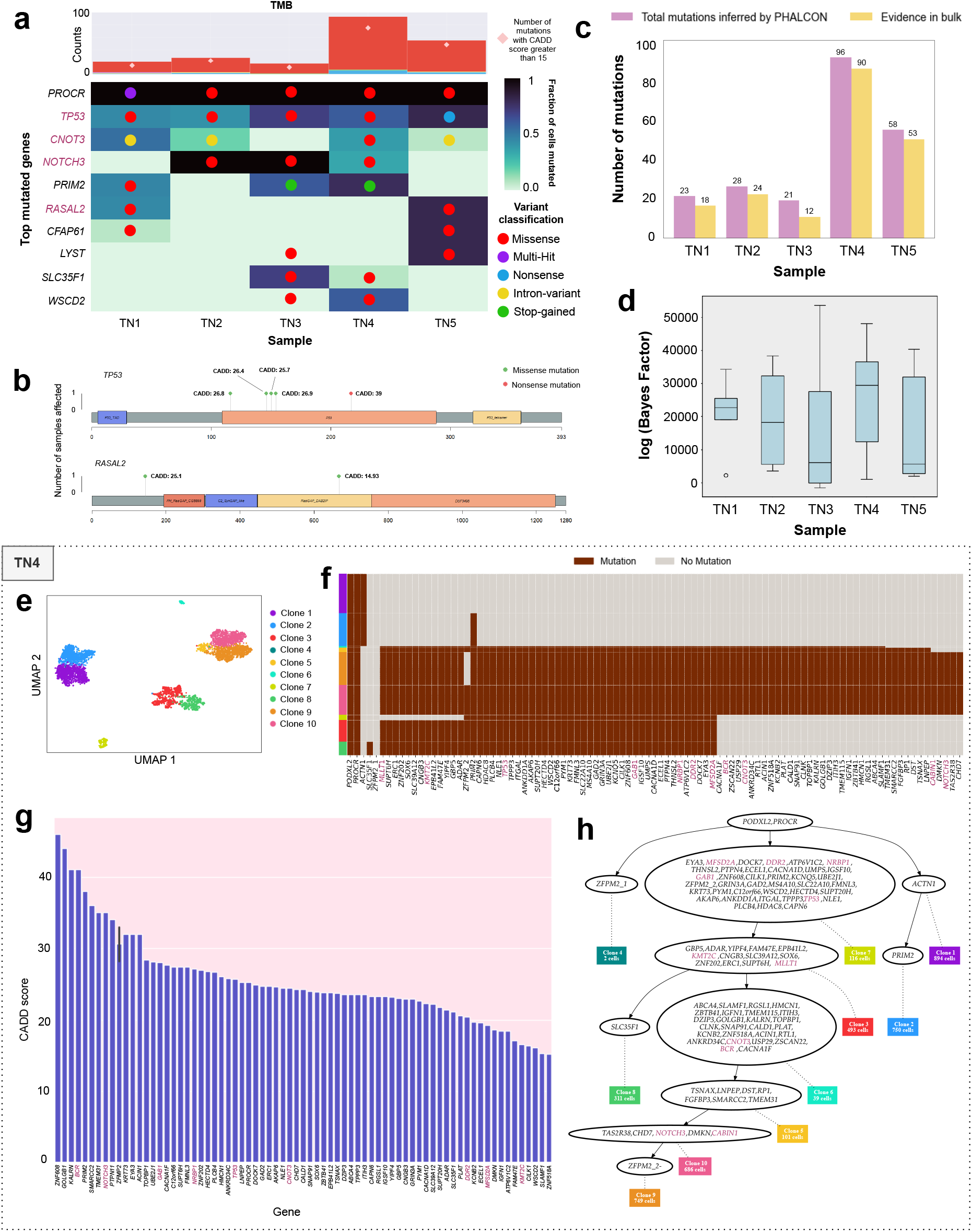
Application of PHALCON on triple negative breast cancer (TNBC) MPT-seq datasets. (a) Fraction of cells harboring a certain mutation for top mutated genes inferred by PHALCON across five TNBC samples. The top part of the figure shows the total mutation burden (TMB) and the number of mutations with CADD score *>* 15 for each TNBC patient. The circles are colored according to the type of the mutation harbored by a gene in a patient. (b) Lollipop plots for *TP53* and *RASAL2* with CADD score annotated mutations. (c) Number of mutations inferred by PHALCON and the number of mutations which had evidence in bulk data for each TNBC patient. (d) Boxplot of log(Bayes Factor) values for the mutations not having evidence in bulk data. The box depicts the median and the interquartile range (IQR, the range between the 25^*th*^ and 75^*th*^ percentile); whiskers extend to 1.5 times the interquartile range; and outliers beyond this range are denoted as empty circles. (e) Visualization of the clones inferred by PHALCON in UMAP for sample TN4. (f) Heatmap of clonally clustered mutations in TN4, as inferred by PHALCON, illustrating the clonal substructure and heterogeneity within tumor subpopulations. The clonal membership of individual cells is represented in the leftmost column. (g) Mutations ranked according to their CADD scores denoting the functional impact of mutations’ deleteriousness in sample TN4 (CADD*>* 15, top 5%). (h) Clonal phylogeny reconstructed by PHALCON delineating the evolutionary history of the sample TN4. The leaves (rectangular boxes colored according to clone identity) represent different sub-clones. Intermediate nodes contain subclonal mutations that are passed down the phylogeny. The root node contains the clonal mutations present across all cells. Oncogenes and Tumor suppressor genes are highlighted in purple across all panels.

We next investigated whether the mutations detected by PHALCON were also detected in the matched bulk exome sequencing dataset from the corresponding patient. Across all five patient samples, a high percentage of mutations (∼ 87.1%) were also detected in the bulk sequencing dataset (Fig. 3c). For the mutations that were detected by PHALCON but were not detected in the bulk data, we performed a statistical test to determine whether these reported mutations can be caused by false positive error (see Methods). For 28 (out of 29) mutations, the statistical test resulted in a large Bayes factor indicating that these mutations cannot be caused by false positive error (Fig. 3d).

### PHALCON delineates mutational substructure of TNBC tumors

PHALCON detected the presence of multiple clones in each TNBC sample and inferred a more intricate clonal evolutionary history as compared to what was reported in the original study. In each TNBC sample, PHALCON resolved more subclonal populations (up to 10 subclones) and detected rare subclones (0.03%-8.84% of the population). For TN4, originally reported to have 3 tumor clones, PHALCON identified a more resolved subclonal landscape consisting of 10 distinct clones, including a rare clone comprising of just 2 cells (0.048% of the population) (Fig. 3e). Clustered heatmap of the mutation profiles in TN4 further revealed two clonal mutations shared by all clones and 91 mutations shared by 7 clones, including mutations in *TP53*, as well as other tumor suppressor genes (*NRBP1, DDR2*) and oncogene (*GAB1*) (Fig. 3f). To predict the functional impact of these mutations, we use Combined Annotation Dependent Depletion (CADD) tool [35], which identified several oncogenes and tumor suppressor genes (e.g. *BCR, NOTCH3, GAB1, NRBP1, TP53*) to harbor mutations with high functional impact scores (Fig. 3g). PHALCON further inferred the clonal phylogeny tree and the order of the mutations during the clonal evolution (Fig. 3h). Truncal mutations that occurred in all clones included the missense mutation in *PROCR*. While three clones (clones 1, 2, and 4) did not expand much in terms of mutations, clone 7 acquired 39 mutations (shared by rest 6 clones as well) including *TP53, GAB1, NRBP1, DDR2* and led to the expansion of the tumor through acquisition of more subclonal mutations which gave rise to newer tumor clones. The divergence of a tumor subclone from the most recent common ancestor in most cases were associated with the gain of mutation in known oncogene or tumor suppressor gene (e.g., *KMT2C* and *MLLT1* for clone 3, *CNOT3* and *BCR* for clone 6, and *NOTCH3* for clones 9 and 10).

Similarly to TN4, TN3 and TN5 also harbored large number of subclones (8 and 9 respectively) (Supplementary Figs. 4a, 4e). While both of them contained a clonal mutation in *PROCR* (Supplementary Figs. 4b, 4f), TN3 also harbored a clonal mutation in *NOTCH3* that was shared by all tumor clones (Supplementary Fig. 4b). In TN3, the mutations with the highest impact scores included mutations in *TP53, PLEKHA6, PRIM2* (Supplementary Fig. 4c), while in TN5, significant mutations were detected in *TP53, OCRL*, and *BAP1*, among other genes (Supplementary Fig. 4g). For both TN3 and TN5, PHALCON inferred a highly branched clonal phylogeny (Supplementary Figs. 4d, 4h). In TN3, 4 clones (clones 1, 2, 3, and 8) emerged from a descendant of the ancestral clone (clone 4 harboring 4 clonal mutations) that acquired 11 sub-clonal mutations (including *TP53*), whereas the other three clones (clones 5, 6, 7) directly descended from the ancestral clone with the addition of only 1 or 2 mutations (Supplementary Fig. 4d). Similarly, for TN5, clone 5 diverged from the ancestral population (clone 6) by acquiring a large set of somatic mutations (including high CADD-score mutations in *TP53, BAP1*, and *NDST4*) and gave rise to five other clones (clones 1, 4, 7, 8, 9) which harbored a few additional mutations (Supplementary Fig. 4h). Two other clones (clones 2 and 3) directly descended from clone 6 with the acquisition of only 1 mutation.

The remaining two patients, TN1 and TN2, harbored a smaller number of clones (6 and 3 respectively) compared to other three patients (Supplementary Figs. 5a, 5e). While both the patients contained a clonal mutation in *PROCR* (Supplementary Figs. 5b, 5f), TN2 also contained a clonal mutation in *NOTCH3* (Supplementary Fig. 5f). In TN1, PHALCON detected high-impact mutations (high CADD scores) in several oncogenes and tumor suppressor genes, including *CDS1, TP53*, and *RASAL2* (Supplementary Fig. 5c). In the clonal phylogeny inferred by PHALCON, two clones (clones 1 and 4) descended from the ancestral population (clone 3 consisting of only 47 cells, 1.1% of the total population), and both then followed a linear evolution with the addition of one or two mutations (Supplementary Fig. 5d). While clone 1 acquired only one mutation (*CNOT3*) in diverging from the ancestral population, the divergence of clone 4 was associated with the acquisition of 18 mutations, including mutations in several oncogenes and tumor suppressor genes (e.g., *TP53, MEFV, CDS1, RASAL2*).

The evolution of TN2, on the other hand, was simpler, harboring three different clones, with the majority of cells (57.83%) belonging to the ancestral clone (clone 1) carrying the clonal mutations in *NOTCH3, PROCR*, and *PODXL2* (Supplementary Fig. 5h). The direct descendant of clone 1 diverged into two subclones, clone 2 and clone 3, both carrying alterations in tumor suppressor genes *CNOT3* and *DICER1* respectively (Supplementary Fig. 5h). Both clone 2 and clone 3 shared 17 mutations from the common ancestor, including high CADD-score mutations in *TP53, NBN*, and *KAT6A* (Supplementary Fig. 5g).

### PHALCON delineates cellular-level mutational landscape in acute myeloid leukemia

Next, we applied PHALCON to a large cohort of AML patients (*n* = 123) for which scDNA-seq was performed on a total of 735,483 bone marrow mononuclear cells (BMMCs) using Mission Bio Tapestri platform [24]. For these samples, scDNA-seq was conducted using two different targeted panels: a panel of 19 genes, and a custom panel covering 37 genes. PHALCON detected 294 (unique) somatic mutations in 34 cancer genes including 195 SNVs and 99 small insertion-deletions (indels). Figure 4a shows the top 20 mutated genes across all 123 patients, where *FLT3* was mutated in the highest number of samples (*N* = 38, 30.8% 27 with ITD and 14 with non-ITD mutations), followed by *NRAS* (*N* = 38, 30.8%), *DNMT3A* (*N* = 36, 29.2%), *NPM1* (*N* = 27, 21.9%), *U2AF1* (*N* = 26, 21.1%), *IDH2* (*N* = 24, 19.5%), *RUNX1* (*N* = 23, 18.6%), *EZH2* (*N* = 20, 16.2%), *ASXL1* (*N* = 18, 14.6%), *TET2* (*N* = 18, 14.6%), *PTPN11* (*N* = 17, 13.8%), and *WT1* (*N* = 17, 13.8%). PHALCON was able to detect somatic mutations in the known driver genes associated with the major pathways for AML (Supplementary Fig. 6). PHALCON further inferred the tumor evolutionary history for these patients and it inferred linear clonal evolution for 64 (52%) patients, branching clonal evolution for other 59 (48%) patients (Fig. 4a). As anticipated, samples with branching evolution harbored larger number of clones compared to those with linear evolution (Supplementary Fig. 7a). Compared to the original study, PHALCON was able to detect more somatic mutations in multiple genes including *U2AF1, BCOR, EZH2, PHF6, STAG2* and *KIT* (Fig. 4b). Moreover, PHALCON detected somatic mutations in genes *NF1* and *ZRSR2* which were not reported to be mutated in the original study (Fig. 4b). We used CADD to predict the functional impact of the mutations detected by PHALCON and most of the previously reported mutations as well as the newly detected mutations were found to have high CADD score indicating potential functional impact of the mutations (Fig. 4c, Supplementary Fig. 7b). Specifically, *U2AF1* was found to harbor a SNV (p.C18G) in its CCCH Zinc finger (Znf) domain in 21 samples, and the SNV was predicted to have a high functional impact by CADD (Fig. 4d). MutPred2 predicted high pathogenicity score for this SNV (score = 0.962) with multiple mechanisms altered (Supplementary Fig. 8a). Similarly, the SNV (p.G291N) present in *PHF6* in 12 samples (Fig. 4e) was also predicted to have a high pathogenicity score by MutPred2 (Supplementary Fig. 8b). Moreover, PHALCON detected missense mutations with support in multiple samples in *BCOR*, a transcriptional repressor known to be involved in hematologic diseases [36] (Fig. 4f). To further investigate whether mutations in these genes (for which PHALCON detected more somatic mutations) could influence the clinical outcomes in AML patients, we integrated and analyzed somatic mutation data from the Oregon Health & Science University (OHSU) BeatAML 2.0 cohort (942 samples from 805 patients) [37] and TCGA AML cohort (200 patients) [38] resulting in a cumulative cohort of 1005 AML patients. In this cumulative cohort, patients harboring mutations in the genes *U2AF1, BCOR, PHF6* and *ZRSR2* were found to have poor overall survival (Fig. 4g, Supplementary Fig. 9a-b).

**Figure 4:**
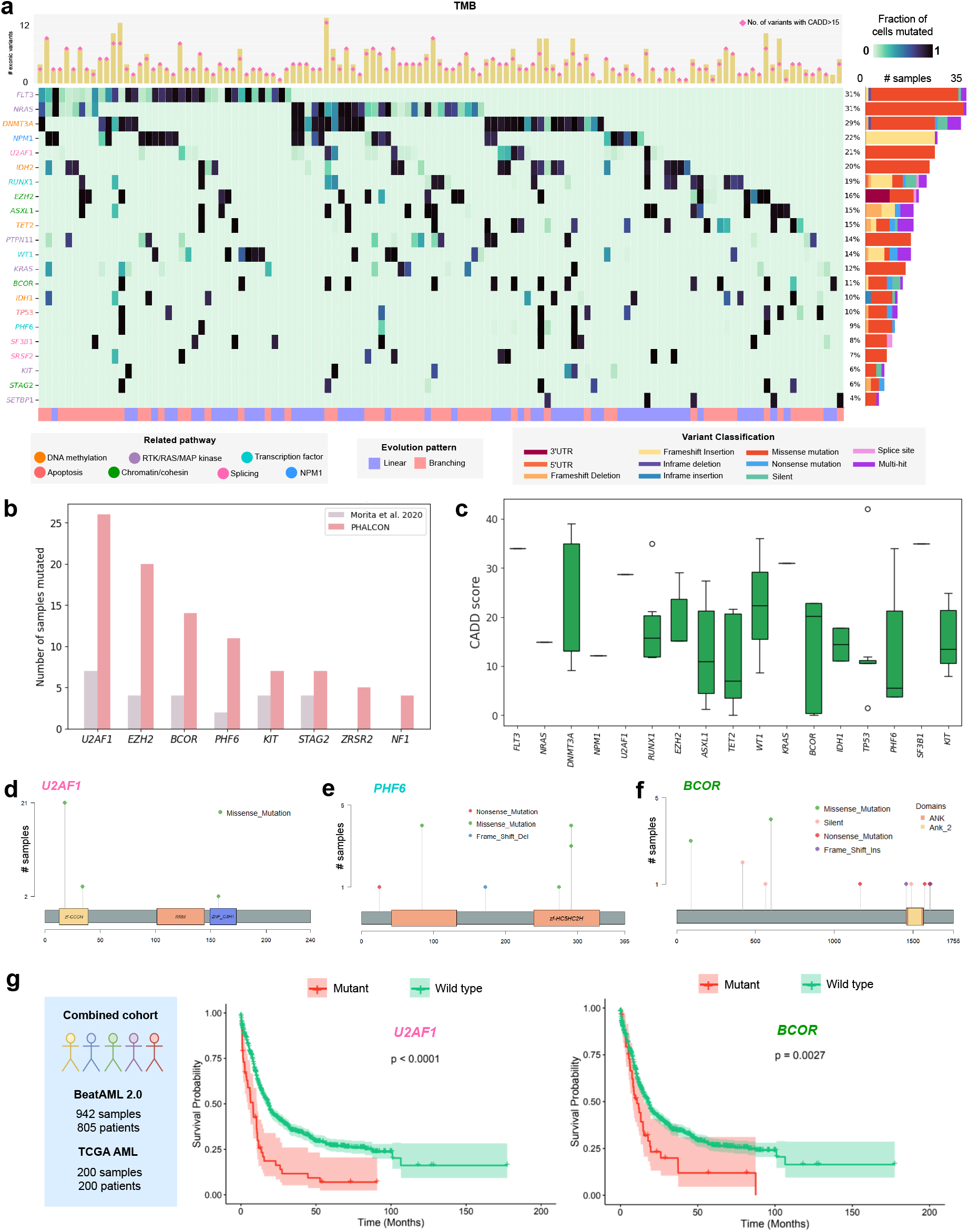
Application of PHALCON on acute myeloid leukemia (AML) datasets. (a) Fraction of cells harboring a certain mutation for top mutated genes inferred by PHALCON across 123 AML samples. The genes are colored according to their associated pathways. The top part of the figure shows the total mutation burden (TMB) and the number of mutations with CADD score *>* 15 for each AML patient. The color strip at the bottom indicates the evolution history pattern (linear/branching) across all samples. The barplot on the right shows the variant classification across all samples for each of the top mutated genes. (b) Comparison of number of mutated samples for genes for which PHALCON detected mutations in more samples as compared to the original analysis. (c) Boxplot of CADD scores for PHALCON-detected variants in top mutated genes, which were not reported in the original study [24]. Lollipop plots for (d) *U2AF1*, (e) *PHF6*, and (f) *BCOR* genes illustrating the mutations detected by PHALCON. (g) Survival curves for *U2AF1* and *BCOR* mutations for a cohort of 1005 patients combined from BeatAML 2.0 [37] and TCGA AML [38].

We also validated the PHALCON-detected mutations using the bulk exome sequencing datasets from the corresponding patients. Matched bulk data was available for 90 patients and considering exonic SNVs and small indels, in each patient, a large fraction (∼ 70.05% on average) of PHALCON reported variants were orthogonally validated by the bulk data (Supplementary Fig. 10a). For the mutations that were reported by PHALCON but not detected in the bulk data, we performed the same statistical test (see Methods) as done for the TNBC datasets to evaluate whether these reported mutations can be caused by false positive errors. For all such mutations, the statistical test resulted in a large Bayes factor indicating that these mutations cannot be caused by false positive error (Supplementary Fig. 10b).

### PHALCON identifies subgroups of AML patients with poor survival characterized by mutations in specific pathways

To understand whether specific mutational profile detected by PHALCON is associated with poor prognosis in AML, for each patient, using the phylogenetic tree inferred by PHALCON we derived a pathway mutation profile based on the pathways affected by the top mutated genes (see Methods for details). The pathways included DNA methylation, receptor tyrosine kinase (RTK)/Ras GTPase (RAS)/MAP Kinase (MAPK) signaling (RTK-RAS), chromatin/cohesin, *NPM1*, transcription factor, splicing and apoptosis [39] (Supplementary Table 2). Graph-based clustering of these pathway mutation profiles resulted in five clusters (pathway subgroups) which had distinct pathway mutation profiles (fraction of cells harboring mutations in specific pathways) (Fig. 5a, Supplementary Fig. 11a). Interestingly, these pathway subgroups varied based on the pattern of tumor evolution as well (Supplementary Fig. 11b), for example, subgroup 1, characterized mainly by high fraction of cells harboring DNA methylation mutations had higher proportion of linear trees (66% linear, 34% branching), whereas for subgroup 4 involving RTK/RAS/MAP kinase-associated mutations, branching evolution was observed in larger fraction of samples (38% linear, 62% branching). Other subgroups had similar proportions of linear and branching trees. All five subgroups were similar in terms of median number of subclones (Supplementary Fig. 11c). To compare the pathway subgroups, we utilized the ELN2022 AML risk classification system [40] by using the subset of genetic abnormality parameters overlapping with our mutated genes (see Table 6 of [40]) to assign the samples to one of the categories: favorable, intermediate, and adverse. Subgroups 1, 2, and 5 had higher proportion of samples assigned to the adverse category (68.2%, 100% and 70% respectively) as compared to subgroups 3 and 4 (Fig. 5b). Moreover, 7.3% samples in subgroup 1 were assigned to the intermediate category and the samples in subgroups 1 and 2 harbored higher number of mutations associated with the adverse risk category (Supplementary Fig. 11d).

**Figure 5:**
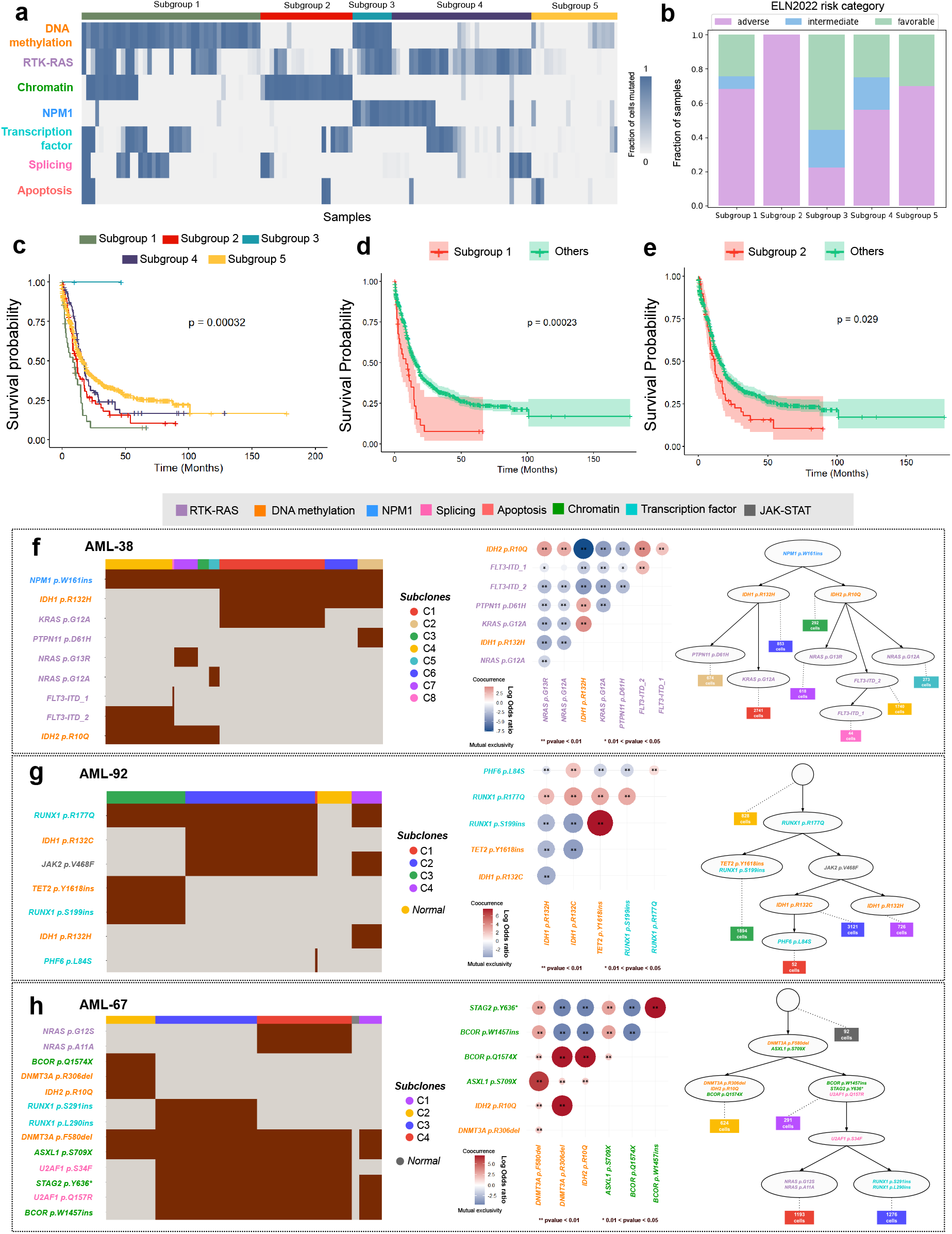
PHALCON uncovers subgroups of patients with poor survival harboring mutations in specific pathways and cellular-level mutual exclusivity and co-occurrence of AML driver mutations. (a) Heatmap showing the fraction of cells harboring mutations in a specific pathway inferred by PHALCON across 123 samples. The samples are clustered into five groups based on the pathway mutation profile. The cluster association of each sample is represented in the topmost row. (b) Distribution of 123 AML samples to one of the three ELN2022 risk categories (adverse, intermediate, favorable) across different subgroups. (c) Survival curves of all five patient subgroups after mapping to the larger combined cohort (TCGA AML and BeatAML 2.0). (d) Survival curve of subgroup 1 patient samples in the combined cohort compared to all other patient samples. (e) Survival curve of subgroup 2 patient samples in the combined cohort compared to all other patient samples. (f-h) Three representative cases showing the cellular-level mutual exclusivity and co-occurrence of genes associated with various pathways in AML samples. (f) Cell-level mutual exclusivity patterns of RTK/RAS/MAP kinase pathway mutations and DNA methylation mutations. (g) Cell-level co-occurrence of DNA methylation (*TET2*) and transcription factor (*RUNX1*) mutations. (h) Cell-level co-occurrence of chromatin/cohesin mutations (*BCOR* and *STAG2*). For (f-h), (left) Heatmap of clonally clustered mutations, illustrating the distribution of the driver mutations across different tumor clones. Each row represents a mutation and each column represents a cell. The clonal membership of individual cells is represented in the topmost row. (middle) Log odds ratio plot depicting the pairwise association between mutations in the patient sample. The color and size of a panel denote the magnitude of the logarithmic odds ratio (log OR). The reference bar maps the colors to their corresponding log OR values. Red denotes co-occurrence, while blue indicates mutual exclusivity. The statistical significance of these associations, based on p-values is marked by asterisks: * 0.01 *<* pvalue *<* 0.05, ** pvalue *<* 0.01. (right) Clonal phylogeny reconstructed by PHALCON showcasing the evolutionary history of the sample. The leaves (rectangular boxes colored according to clone identity) represent different subclones. Intermediate nodes contain sub-clonal mutations that are passed down the phylogeny. The root node contains the clonal mutations present across all cells. Genes are colored according to their associated pathways across all three panels.

Given the distinct pathway mutation profile and ELN2022 risk profile of the five subgroups in the single-cell AML cohort, we mapped them to the larger cohort of 1005 samples combining TCGA AML and BeatAML 2.0, where each sample in the combined cohort was mapped to one of these five subgroups based on the similarity of its pathway mutation profile to the average pathway mutation profile of the subgroup (see Methods). Survival analysis of these 5 subgroups in the larger combined cohort showed significant difference in survival across all the 5 subgroups (p-value = 0.00032, Fig. 5c). Subgroup-wise survival analysis further showed that subgroup 1, characterized by the presence of DNA-methylation mutations in high fraction of cells had poor overall survival (p-value = 0.00023, Fig. 5d). Similarly, subgroup 2 characterized by chromatin/cohesin mutations in a high fraction of cells also showed poor survival as opposed to other groups (p-value = 0.029, Fig. 5e). Subgroup 3 was characterized by the presence of DNA methylation, *NPM1*, and RTK/RAS/MAP kinase mutations and only 3 samples in the combined cohort were mapped to this subgroup. Subgroup 4 was characterized by the presence of RTK/RAS/MAP kinase mutations in a high fraction of cells and it was associated with poor survival but the effect was not statistically significant (Supplementary Figs. 11e-f). Thus, PHALCON was able to characterize subgroups of AML patients in terms of mutations in major pathways and also identified specific subgroups with poor prognosis.

### PHALCON reveals clonal relationships of oncogenic mutations in acute myeloid leukemia

PHALCON was further able to identify co-occurrence and mutual exclusivity among driver mutations at cellular resolution. Several AML patients harbored multiple mutations in different genes (*FLT3, NRAS, KRAS, PTPN11*) involved in receptor tyrosine kinase (RTK)/Ras GTPase (RAS)/MAP Kinase (MAPK) signaling pathway and these mutations were mostly subclonal and observed to be present in mutually exclusive clones at the cellular level. As reported in the original study, similar mutual exclusivity was observed for genes (*DNMT3A, TET2, IDH1*, and *IDH2*) involved in DNA methylation. For example, in AML-38 (Fig. 5f), an early mutation in *NPM1* provided a selective advantage to DNA methylation mutations, leading to the divergence of two independent clones harboring *IDH1* p.R132H and *IDH2* p.R10Q mutations. Subsequently, these two branches diverged into two independent subclones harboring *PTPN11* p.D61H and *KRAS* p.G12A, and three independent subclones harboring *NRAS* p.G12A, *NRAS* p.G13R and *FLT3-ITD*, respectively. The log odds ratio plot from the pairwise association analysis demonstrates this cellular mutual exclusivity among RTK-RAS-associated mutations and DNA methylation mutations, with almost all associations having significant p-values. Other examples of RTK-RAS-MAPK and DNA methylation mutations exhibiting mutual exclusivity are presented in Supplementary Figs. 12 and 13 respectively. In line with the original study [24], PHALCON also found *TP53* and *PPM1D* mutations to be mutually exclusive. In three different samples that harbored mutations in *PPM1D, PPM1D* and *TP53* mutations occurred in mutually exclusive subclones (Supplementary Fig. 14). In addition, PHALCON uncovered some novel cellular co-occurrences of driver mutations from these AML samples. In multiple patient samples, PHALCON identified DNA methylation mutations and variants in transcription factors to co-occur on the phylogeny. One such sample, AML-92 (Fig. 5g), shows small insertions in *TET2* p.Y1618ins and *RUNX1* p.S199ins to co-occur in one of the clones. The same sample also exhibited the mutual exclusivity of two DNA methylation mutations, *IDH1* p.R132C and *IDH1* p.R132H. Other samples such as AML-99, AML-104, and AML-105 (Supplementary Fig. 15a-c) also demonstrate the co-occurrence of *DNMT3A*-*RUNX1, TET2 PHF6*, and *DNMT3A*-*RUNX1* respectively. Some of these samples simultaneously demonstrated the cellular mutual exclusivity of DNA methylation mutations. The cellular-level co-occurrence of chromatin-cohesin mutations was also observed in several samples. Sample AML-67 (Fig. 5h), for instance, shows a stopgain mutation *STAG2* p.Y636* and a small insertion *BCOR* p.W1457ins to co-occur in one of the two clones emerging from another chromatin-associated mutation *ASXL1* p.S709X. Other samples harboring such co-occurrences are presented in Supplementary Fig. 16.

## Discussion

Single-cell DNA sequencing (SCS) provides unprecedented insights into somatic cell evolution by enabling the direct investigation of genetic variability between individual cells. SCS is particularly promising for cancer biology, as intra-tumor heterogeneity (ITH) where multiple distinct tumor cell populations co-exist can lead to therapeutic resistance and cancer relapse. As a result, it is necessary to characterize ITH by elucidating the mutational composition of the cell populations as well as the chronological order of the mutations. Recently developed high-throughput amplicon-based panel sequencing platforms (e.g., Tapestri by Mission Bio) aim to resolve ITH by profiling the DNA of thousands of cells by targeting a panel of disease-specific genes. However, existing mutation calling methods cannot handle such large number of cells. Here, we introduced PHALCON, the first scalable single-cell variant caller that can simultaneously perform variant calling and reconstruct the clonal substructure and phylogenetic history of a tumor sample from single-cell panel sequencing datasets consisting of thousands of cells. PHALCON employs a finite-sites model of evolution to permit recurrent mutations on the tumor phylogeny and can infer somatic SNVs and small indels from the read count data of thousands of cells generated using panel sequencing platform. PHALCON employs graph-based clustering for inferring the clonal substructure of a tumor and a maximum-likelihood-based framework for inferring the clonal phylogeny and the best possible mutation history of the genomic sites which in turn help to finalize the variants and genotype them in individual cells.

We benchmarked the variant calling performance of PHALCON against that of state-of-the-art SCS variant callers SCIPhIN [22], Phylovar [20] and SIEVE [23] using a comprehensive set of simulated datasets encompassing a wide variety of experimental settings. PHALCON had near-perfect variant calling for all these datasets (F1 score ≈ 0.99). In comparison, SIEVE failed to produce any result for these datasets, SCIPhIN and Phylovar could only handle the smallest datasets (2000 cells) and failed to produce any result for larger datasets (5000 and 10000 cells). PHALCON substantially outperformed the other variant callers across all experimental settings. Even for datasets harboring copy number alterations which are not modeled by PHALCON, PHALCON retained its superior performance across varying levels of copy number alterations while other methods’ performance monotonically degraded with the increase of copy number alterations. PHALCON further exhibited a massive improvement (44 − 47.5×) in runtime over SCIPhIN and Phylovar. PHALCON’s runtime also increased linearly with the number of cells making it suitable for handling such large datasets.

We further evaluated PHALCON’s ability to reconstruct the evolutionary history of the cells against that of SCIPhIN, Phylovar, and COMPASS [29], a state-of-the-art phylogeny inference method capable of handling panel sequencing datasets. Even though ground truth mutation sites were passed as input to COMPASS, it ended up removing multiple true mutation sites and failed to capture the underlying evolutionary history of the cells. Across all simulated datasets across all experimental settings, PHALCON outperformed all other methods in reconstructing the phylogenetic tree by large margins (410.43 − 32931.78% improvement over other methods).

We applied PHALCON on panel sequencing datasets from five triple negative breast cancer (TNBC) patients for which it detected novel somatic mutations with high functional impact. Particularly, PHALCON inferred a novel clonal mutation in *PROCR* gene in all five patients and this mutation was predicted to have high functional impact and pathogenicity. Since *PROCR* has been shown to enrich for Wnt1 basal-like tumors and its inhibition has been found to result in decrease in tumor growth [32], we further investigated *PROCR*’s expression profile in TNBCs and found higher expression of *PROCR* in cancer epithelial cells as compared to normal epithelial cells in TNBC patient and in TCGA cohort, higher expression of *PROCR* was associated with poor survival. Thus PHALCON was able to identify novel somatic mutation in a gene important for TNBC progression. For all five patients, PHALCON was able to resolve the cellular subpopulations in much higher resolution by uncovering more tumor clones as compared to the original study and also identified rare clones as small as consisting of only two cells. Moreover, PHALCON was able to provide a more comprehensive understanding of the mutational order of the clonal evolution where it identified important oncogenes and tumor suppressor genes to harbor deleterious mutations at the major lineages of the clonal phylogenies.

Next, we analyzed a large acute myeloid leukemia (AML) panel sequencing dataset (123 patients) using PHALCON. Compared to the original study, PHALCON reported more somatic mutations with high functional impact scores in several genes (e.g., *U2AF1, BCOR, PHF6*) that are reported to harbor driver mutations in other AML cohorts [41]. Specifically, PHALCON revealed a new recurrent variant of high pathogenicity in *U2AF1* in its zinc finger domain, a known hotspot in splicing factor mutations associated with adverse outcomes [42, 43], mutations in *PHF6*, a chromatin regulator known to harbor loss of function mutations in AML [44], and BCOR alterations, which are recurrent in myeloid malignancies and linked to transcriptional dysregulation [45]. Moreover, PHALCON also detected mutations in genes that were completely missed by the original study: *ZRSR2*, a component of the minor spliceosome known to be recurrently mutated in AML cases with splicing factor mutations [42] and *NF1*, a RAS-MAPK-signaling pathway gene recurrently mutated in AML [46]. Our results further demonstrate that pathway-based clustering, informed by PHALCON detected single-cell mutation profiles, captures heterogeneity in AML evolution and provides insights into the impact of specific mutational patterns on disease progression. Notably, two patient subgroups characterized by high fractions of DNA methylation and chromatin/cohesin mutations, respectively, exhibited significantly worse overall survival and this finding is consistent with previous studies showing the critical role of DNA methylation dysregulation in AML progression [47] and poor prognosis of AML patients harboring cohesin mutations [41]. The ability of PHALCON to identify co-occurrence and mutual exclusivity among driver mutations at cellular resolution provided deeper insights into the clonal architecture and evolutionary dynamics of AML. PHALCON reported mutations in key signaling pathways, particularly RTK-RAS-MAPK and DNA methylation, to frequently occur in mutually exclusive subclones, reinforcing prior observations of functional redundancy and selective pressures shaping leukemic evolution [41, 24]. Moreover, PHALCON highlighted the cellular co-occurrence of DNA methylation mutations with transcription factor mutations which were reported to co-occur in the same patient by earlier studies [41, 48]. PHALCON further uncovered the cellular co-occurrence of chromatin/cohesin mutations suggesting that such alterations may contribute to clonal expansion by promoting transcriptional reprogramming and synergistically perturbing chromatin looping, which have previously been implicated in AML pathogenesis [49].

PHALCON represents a major leap forward for variant calling in single-cell panel sequencing datasets. By harmonizing finite-sites evolutionary modeling with a scalable joint inference framework, PHALCON is the only phylogeny-aware single-cell variant caller which can perform improved variant calling and reconstruct tumor phylogenies harboring complex mutational events across thousands of cells. By doing so, it unlocks the potential of high-throughput single-cell genomics to dissect ITH, trace clonal evolution, and identify clinically actionable mutations. Given the increasing size and wider applicability of the single-cell panel sequencing datasets, we envision PHALCON to play an important role in mutation calling and clonal lineage inference from these datasets for translating cellular-resolution data into insights for cancer biology and therapeutic development.

## Methods

### Overview

PHALCON first identifies the putative mutation loci, utilizes the putative sites for inferring clonal populations and then employs a maximum likelihood-based algorithm for simultaneously performing variant calling and clonal phylogeny inference. We first describe the model for calculating mutation likelihood for each cell at each genomic site. After that, we provide a detailed description of the steps involved in PHALCON.

### Computation of mutation likelihoods

PHALCON employs a likelihood-based approach in determining the presence of a mutation at a genomic locus (site) of a cell. Assuming *a*_*ij*_ and *d*_*ij*_ to be the alternate allele count and total read depth respectively at the *i*^*th*^ site of *j*^*th*^ cell, we compute the mutation likelihood of a site *i* in cell *j*, using genotype likelihoods for each possible genotype. Following previous single-cell phylogeny methods [15, 16], we consider three possible genotypes at a site - homozygous reference, heterozygous, and homozygous alternate. To compute the genotype likelihoods, we adopt the alternate read distributions from Lodato et al. [50], which account for skewed read counts.

The likelihood for a homozygous reference genotype *l*(*g* = 0) is modeled using a non-symmetric beta-binomial distribution with parameters *α*_*q*_ and *β*_*q*_:

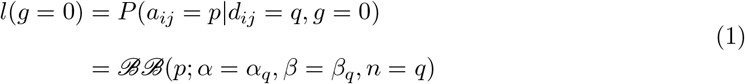

Homozygous alternate genotype may appear due to the deletion of the reference allele at a site containing a mutation. This event may cause the read counts to only contain the evidence of the alternate allele. To compute the likelihood of the homozygous alternate genotype, we simply switch the parameters of the beta-binomial distribution as mentioned in the homozygous reference case. The likelihood of a homozygous alternate genotype *l*(*g* = 1) is calculated as:

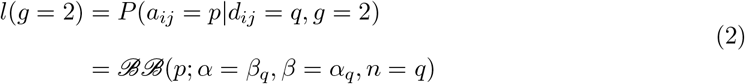

A major drawback of single-cell sequencing is the imbalanced amplification of the two alleles. For a heterozygous genotype, conventional beta-binomial distribution might not properly capture the imbalanced amplification. Lodato et al. [50] empirically showed that the alternate depth distribution of the confident heterozygous sites has a bell-shaped curve with sharp peaks at the tails. Following this, we model the alternate read distribution for a heterozygous genotype using a mixture of two symmetrical beta-binomial distributions as given by:

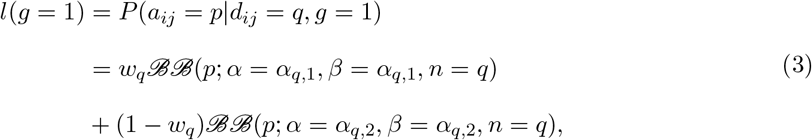

where *α*_*q*,1_ and *α*_*q*,2_ represent the parameters of the two symmetric beta-binomial distributions that the mixture consists of and *w*_*q*_ represents the mixture parameter representing the allelic dropout rate. Supplementary Fig. 17 shows how the mixture of beta-binomial distribution fits the alternate read counts for different values of total read depth for real tumor datasets.

In equations (1) to (3), ℬℬ denotes beta-binomial distribution whose parameters scale linearly as the total read depth (*q*) increases. Similarly, the mixture model parameters in equation (3) scale up linearly as the read depth increases. These linear relationships are formulated as:

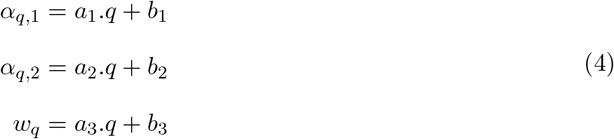

The slopes and intercepts for *α*_*q*_ and *β*_*q*_ for the beta-binomial distributions for homozygous genotypes are directly adopted from [50], whereas the slopes and intercepts (*a*_1_, *a*_2_, *a*_3_, *b*_1_, *b*_2_, *b*_3_) for the mixture model parameters in equation (4) are estimated for individual datasets using the maximum likelihood approach from [50]. Using the likelihoods of genotypes 0, 1, and 2 we calculate the likelihood of mutation *ℓ*_*ij*_ at each site *i* in each cell *j* using normalization as given by:

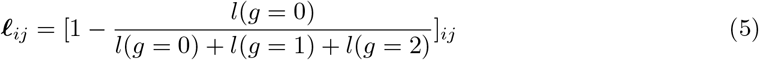

### Identification of putative mutation sites

PHALCON begins with the identification of sites likely to harbor mutations by filtering uninformative sites through a simple five-stage filtering process. The first optional filter removes the read count data at a site with a low genotype quality (GQ) score. Although GQ scores are usually available in the loom files generated by Mission Bio Tapestri, this filter can be turned off if GQ scores are not available. Using the second filter, we remove read count information from the cells with coverage depth less than a given threshold. Following these two filtering steps, PHALCON calculates the mutation likelihood matrix, as described in the earlier section, which serves as the basis for the subsequent filter. The third step filters out read-count information based on low variant allele frequency (VAF) as a VAF value much lower than 0.5 (for a heterozygous variant, expected VAF ∼ 0.5) can indicate a potential false positive. This filter considers cells to harbor a putative mutation, so we only consider cells whose mutation likelihood value is *>* 0.5. After applying three cell-level filters, the fourth filter removes genomic sites where read count information is available for fewer than a threshold proportion of cells, as without sufficient number of cells with high-quality read count information, mutations at a site cannot be confidently determined. Finally, the last filter removes a genomic site if the fraction of cells harboring a mutation is very low. This eliminates any chances of calling a false positive mutation. A detailed description of the filters, along with the default threshold values, can be found in Supplementary Note 1.

### Inference of tumor clones using graph-based clustering

After identifying the putative mutation sites, we determine the cell populations (aka clones) in the tumor sample by employing a graph-based clustering algorithm. We specifically utilize graph-based clustering as it does not pre-assume any cluster structure, making it better suited for capturing heterogeneity. Particularly, we employ spectral clustering for inferring the cell clusters whose optimal number is determined based on the eigen gap heuristic. Using the mutation likelihood vectors (the rows of the matrix ℒ) of *m* single cells, PHALCON constructs an *m* × *m* distance matrix based on Euclidean distance between pairs of cells. PHALCON then constructs an affinity graph based on the pairwise distance matrix using the Radial Basis Function (RBF) kernel and subsequently computes the symmetrically normalized Laplacian matrix *L*_*norm*_ based on which it performs spectral clustering. PHALCON uses the eigen-gap heuristic to determine the optimal number of clusters *K*. For some of the complex real datasets, where spectral clustering reports large clusters and cannot subdivide them into distinct clones, we applied Leiden clustering [51] on each of the large clusters. Leiden clustering optimizes modularity and helps in detecting groups of nodes that are more densely connected than the rest of the network. This helped us recognize more rare cell populations in real tumor datasets. While the default clustering approach used by PHALCON is spectral clustering, users have the option of employing other graph-based clustering including Leiden or Louvain for identifying the cell populations.

### Clonal phylogeny likelihood

Following single-cell phylogeny inference methods [15, 16], PHALCON employs a finite-sites model of evolution for inferring the clonal phylogeny that connects all tumor clones. PHALCON’s finite-sites model considers three possible events on the phylogeny: heterozygous mutation, parallel mutation, and back mutation (i.e., mutation loss). PHALCON’s phylogeny inference model aims to infer the clonal tree topology 𝒯 and the mutation placements *σ* on the branches of the clonal tree. PHALCON models the tumor phylogeny using a clonal tree (a binary tree) consisting of *K* leaves corresponding to the *K* clones inferred by PHALCON. The mutations are placed along the branches of this clonal tree. The heterozygous mutations occur only once in the phylogeny and thus can be placed only on one branch of the phylogeny. In contrast, parallel mutations and mutation losses can occur more than once. For parallel mutations and mutation losses, we consider that the genomic site gets mutated twice on the phylogeny and thus can be realized by placing the mutation on two branches of the phylogeny. For parallel mutations, we consider the placement of the mutation on two independent lineages, whereas for the mutation loss, the second placement must occur in the subtree rooted at the branch where the mutation is first placed. More than two occurrences are not considered given the rarity of such events [21].

Given the observed data consisting of alternate allele count matrix 𝒜 and depth matrix 𝒟 over *n* sites and *m* cells, PHALCON aims to compute the parameters 𝒯 and *σ* that maximize the data log-likelihood log *P* (𝒜, 𝒟|𝒯, *σ*) as given by:

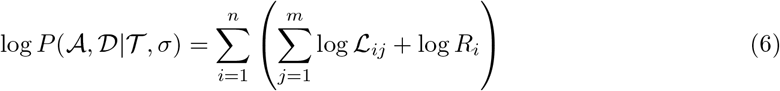

where ℒ_*ij*_ denotes the likelihood of the data at site *i* in cell *j* and *R*_*i*_ is the regularizer corresponding to site *i*. Given the lower frequency of parallel and back mutations, the regularizer is added to penalize the number of occurrences of recurrent mutations to avoid overfitting. This is particularly important as technical artefacts in single-cell sequencing data can resemble the effects of recurrent mutations [21, 22]. *σ* is a vector denoting the mutation attachment to the branches of 𝒯 . For heterozygous mutations, *σ*_*i*_ denotes the branch on which mutation *i* is placed on 𝒯, whereas for the recurrent mutations (parallel and back), *σ*_*i*_ denotes a pair of branches on which mutation *i* is placed on 𝒯 .

The tree topology 𝒯 and mutation placement vector *σ* together give rise to the clonal genotype matrix, *G* = [*g*_*ki*_]_*K*×*n*_, where *g*_*ki*_ represents the genotype of *k*^*th*^ clone at *i*^*th*^ mutation site. *c*_*j*_ denotes the clonal association of cell *j* (*c*_*j*_ ∈ [1, …, *K*]). The genotype of cell *j* at site *i* can be inferred using the clonal association of cell *j* and clonal genotype 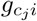 and ℒ_*ij*_ can now be defined as:

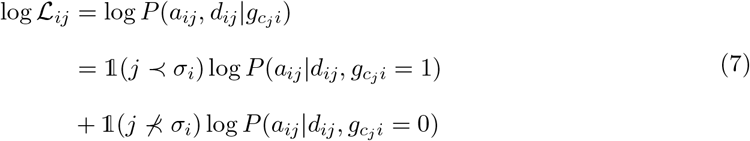

where, the indicator function 𝟙 takes value 1 for *j* ≺ *σ*_*i*_ which denotes the case where clone *c*_*j*_ (and hence cell *j*) in the tree 𝒯 falls below the attachment point (*σ*_*i*_) of mutation *i* (thus cell *j* should harbor mutation at site *i*), and otherwise takes value 0 (genotype of cell *j* is homozygous reference). The terms 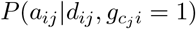 and 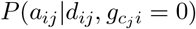 can be obtained from the mutation likelihood matrix ℒ as

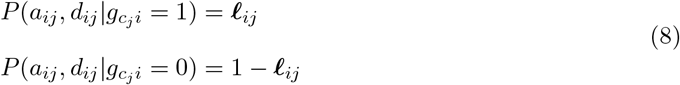

The regularizer *R*_*i*_ for site *i* is defined as:

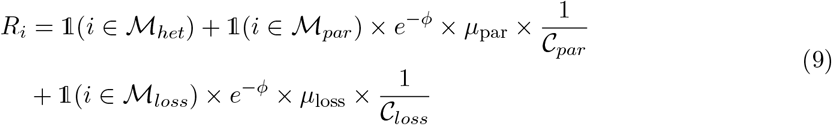

In equation (9), 1 is the indicator function that indicates whether a mutation is inferred to be heterozygous, parallel, or back. ℳ_*het*_, ℳ_*par*_, and ℳ_*loss*_ denote the set of sites harboring heterozygous mutation, parallel mutation, and mutation losses respectively. *ϕ* denotes the regularization coefficient which we set to a large number (700 in our case). *µ*_*par*_ and *µ*_*loss*_ denote the relative frequency of the parallel and back mutation on the phylogeny (set to 0.001 in our experiments). 𝒞_*par*_ and 𝒞_*loss*_ denote the possible number of parallel mutation and back mutation cases on the phylogeny respectively (each case refers to a combination of branches that harbor the mutation) and thus are dependent on *K* (𝒪(*K*^2^)) (see Supplementary Note 2 for examples).

### Inference of clonal phylogeny and mutation placement

To infer the clonal phylogeny and mutation placement that optimize the log-likelihood in equation (6), PHALCON employs a hill-climbing approach similar to that in [20, 15]. At each iteration, a new tree topology is proposed for which the mutations are optimally placed, and the log-likelihood function is evaluated and updated accordingly. We first reconstruct an initial tree topology, 𝒯 ^(1)^ with *K* clones as leaves using the neighbor-joining algorithm [52] which requires pairwise distances between the clones. To calculate the pairwise distances, we first construct an initial clonal genotype matrix, *G*^(0)^, as follows:

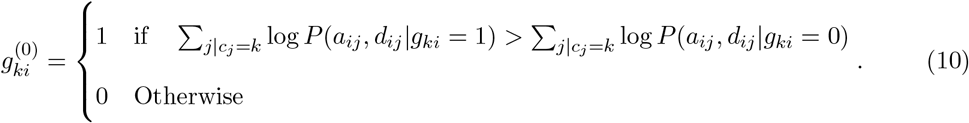

After constructing the initial topology, 𝒯 ^(1)^, PHALCON infers the best mutation history (denoted as 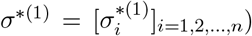) with highest log-likelihood for each site *i* (explained in details in the next section) which further yields the clonal genotype matrix at first iteration, *G*^(1)^ and using equation (6), first log-likelihood value log *P* (𝒜, 𝒟|𝒯 ^(1)^, *σ*^*(1)^) is computed. At each iteration ι *>* 1, tree rearrangement moves [15] including nearest-neighbor interchange (NNI), subtree pruning and regrafting (SPR), and subtree swapping (STS) are employed to propose a new tree topology for which the best mutation history is inferred. The proposed tree and mutation history are accepted if they improve the log-likelihood value from the previous iteration. We also adopt stochastic hill-climbing strategy where the new tree and mutation history are accepted according to the acceptance probability computed as:

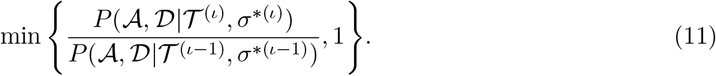

We terminate the search process when the log-likelihood does not improve for a user-defined number of iterations or the maximum number of iterations are exceeded. The search process returns 𝒯 ^*best*^, *σ*^*best*^ that maximize the log-likelihood:

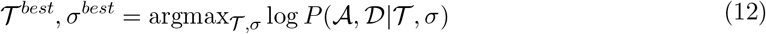

### Inferring the best mutation history on the clonal phylogeny

For a given clonal tree topology at ι^*th*^ iteration, 𝒯 ^(ι)^, each possible mutation history for site *i* leads to a unique genotype configuration at the single-cell level. For each site *i*, we aim to find the mutation history that maximizes the log-likelihood in equation (6). Corresponding to three possible events (heterozygous mutation, parallel mutation, and back mutation) considered by PHALCON’s finite-sites model at a site, we have to consider three types of mutation histories. In this section, we explain how the best mutation history is inferred for site *i* and the notation for site *i* is dropped for the sake of simplicity. For a topology with *K* clones (where leaves are named from {1, 2, …, *K*} for simplicity), let 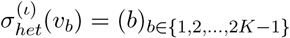 denotes a mutation history where the heterozygous mutation occurs on the branch *b*. Similarly, 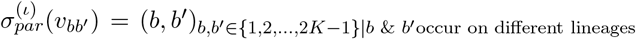 denotes a mutation history due to parallel mutation where the same mutation occurs on branches *b* and *b*^′^.

Let 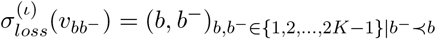 denotes a mutation history corresponding to mutation loss where the first mutation occurs on branch *b* and the mutation gets lost on branch *b*^−^ (*b*^−^ ≺ *b* denotes that *b*^−^ falls below the branch *b* in 𝒯 ^(ι)^). All possible mutation histories due to parallel and back mutation placements in a phylogeny are explained in detail in Supplementary Note 2 (see Supplementary Fig. 18 for examples). Assuming the total number of possible mutation histories under the finite-sites model to be *η* − 1, let 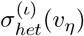 represents the case of no mutation i.e. when none of the branches are mutated. To deduce the effect of all possible mutation histories on the clonal genotype configuration at ι^*th*^ iteration, we define 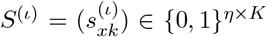 where each row of *S*^(ι)^ represents the genotype configuration corresponding to a particular case from the space of possible mutation histories 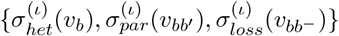.For defining the mapping from mutation histories to the clonal genotypes, we introduce a function *ψ* as follows:

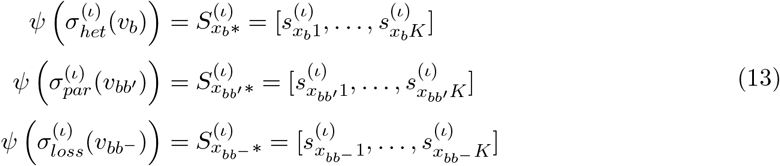

For heterozygous mutation and no mutation, 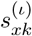 defined as:

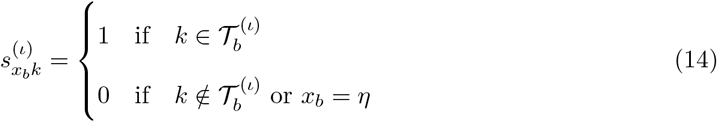

For parallel mutations,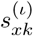 is defined as:

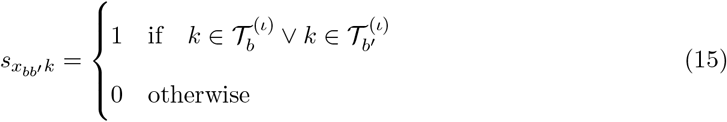

For mutation loss, 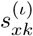 is defined as:

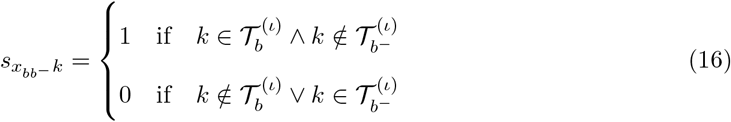

where *k* ∈ {1, …, *K*}, *b, b*^′^, *b*^−^ denote branches in 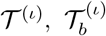 denotes the subtree rooted at branch *b*, and for a leaf 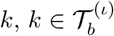 denotes that the leaf *k* occurs in the subtree 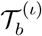,and 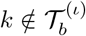 denotes that the leaf *k* does not occur in the subtree 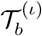.

For the computation of the total likelihood of each possible mutation history for each site across all clones, in addition to *S*^(ι)^, we define two more matrices, *Z*, and *Y* . *Z* is an *m* × *n* matrix that stores the log-likelihoods of no mutation (calculated according to equation (8)), i.e., 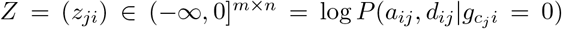.Similarly, transpose of mutation likelihood matrix (ℒ), ℒ ^*T*^ is an *m* × *n* matrix that stores the log-likelihoods of mutation. *Y* is a *K* × *m* clone to cell association matrix which is calculated as:

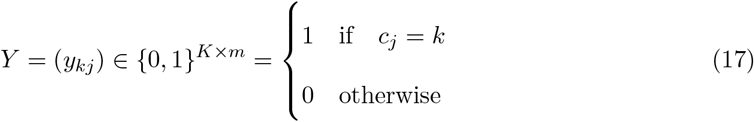

Using the matrices ℒ ^*T*^, *Z, Y*, and *S*^(ι)^, we can calculate the genotype configurations log-likelihood matrix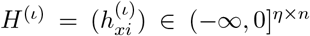,whose each element 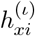 denotes for site *i*, the log-likelihood value of a mutation history corresponding to row *x* of *S*^(ι)^ as shown in equation (13):

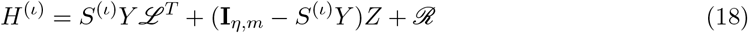

where **I**_*η*,*m*_ is an all-one matrix, ℛ is the regulariser matrix, where each column contains the log-prior regularizer values for all *η* possible mutation histories for site *i* calculated based on equation (9). The best mutation history for each site *i* is found by calculating the index associated with the highest log-likelihood value in the *i*^*th*^ column of *H*^(ι)^:

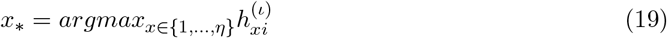

where *x*_*_ ∈ {1, …, *η*} corresponds to the mutation history 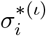 where

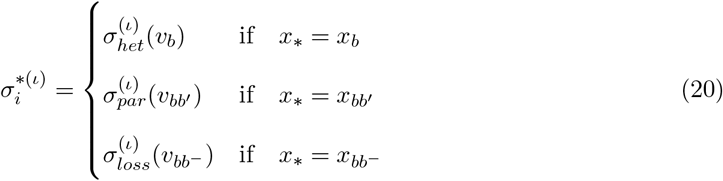

It is important to note that in some cases, a mutation history due to parallel mutation and another mutation history due to mutation loss may lead to the same clonal genotype configuration and in such cases, randomly one mutation history is chosen. Finally, column *i* of the clonal genotype matrix *G*^(ι)^ can be inferred based on 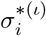:

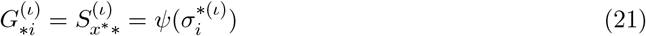

and *G*^(ι)^ is the concatenation of best genotype configurations at all sites:

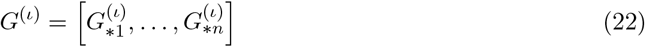

### Inferring the cell genotypes

After finding the maximum likelihood clonal phylogeny 𝒯 ^*best*^ and the best mutation history, *σ*^*best*^, the clonal genotype matrix *G*^*best*^ can be inferred using equations (21) and (22). Based on the cluster associations *c*_*j*_ for cell *j*, where *j* ∈ 1, 2, …, *m*, and *G*^*best*^, the genotypes of individual cells (*G*_*j*_) can be determined as:

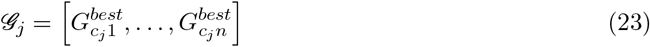

### Simulation of synthetic ground truth datasets

To benchmark the performance of PHALCON against that of other methods, we simulated tumor evolution using a clonal phylogeny and simulated read counts for all single cells across all genomic sites by mimicking the real datasets. We start by simulating a clonal phylogeny with *K* clones by constructing a random binary tree with *K* leaves using Rémy’s Algorithm [53]. Next, we simulate the clonal populations using the simulation strategies from [54, 16], where the clonal prevalences **Φ** = {Φ_1_, Φ_2_, Φ_3_, …, Φ_*K*_} for each clone *k* are sampled from a beta distribution. Considering *m* to be the number of cells that need to be simulated for the dataset, we sample *m* cells from a multinomial distribution with parameters **Φ** as *e*_1_, *e*_2_, …, *e*_*K*_ ∼ *Mult*(*ϕ*). Here, *e*_*k*_ denotes the number of cells sampled from clone *k* and 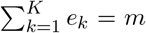.The true genotype of each cell sampled from clone *k* matches the clonal genotype of clone *k*.

Next, we simulate the genotypes for each cell at each site by adopting the strategy from [16]. Assuming *n* ground truth mutation sites, we select a fraction *γ*_clonal_ of the *n* mutations to be clonal, i.e., these mutations occur in all the clones of the tree, and as a result, they are placed at the root of the phylogeny and are populated with heterozygous genotype (*g* = 1). For all other sites, the root node of the tree is populated with homozygous reference genotype (*g* = 0). As we are assuming a finite-sites model of evolution, we allow a small fraction, *γ*_clonal loss_ of the clonal mutations to be lost in the phylogeny. For the remaining sites, we first select the site to harbor either a heterozygous, or a parallel mutation, or a back mutation based on probabilities *p*_*het*_, *p*_*par*_ and *p*_*back*_ respectively. For each mutation type (heterozygous, parallel or back), a particular configuration is selected uniformly randomly e.g., if a particular site is selected to harbor a heterozygous mutation based on the probability *p*_*het*_, then one possible configuration (each configuration corresponds to placing the mutation on a branch excluding the root) for the heterozygous mutation is selected uniformly randomly. Similarly for the parallel and back mutation cases, with uniform probability, we select one mutation history out of all available mutation histories (see Supplementary Note 2). While simulating the datasets, we also ensure that each branch of the phylogeny harbors at least one mutation. For the datasets simulated under infinite sites model, only heterozygous mutations are simulated. The values of the different parameters (*γ*_clonal_, *γ*_clonal loss_, *p*_*het*_, *p*_*par*_ and *p*_*back*_) across different experimental settings are detailed in Supplementary Table 1.

To mimic the errors pertaining to scDNA-seq technologies, we introduced dropouts, amplification errors, and sequencing errors. Once the clonal genotypes are simulated, we can define single-cell genotypes as the genotype of the clone it belongs to. Using a specific dropout probability *µ*_*ADO*_, a site in a cell is selected to be affected by dropout. For such sites, we uniformly drop one of the alleles, i.e., with a probability of 0.5, the genotype is turned into wild type, and for the remaining fraction, the genotype is turned into a homozygous alternate. Using the dropout-affected cellular genotypes, we generated read counts for each cell at each site using the Pólya urn model, similarly as described in [18], with a few modifications to account for the characteristics of the amplicon sequencing datasets. We simulated an artificial reference chromosome with 40000 base pairs, divided this chromosome into shorter chunks of pre-defined length, and sampled coverage values for individual sites from an empirically determined discrete probability distribution using the coverage values extracted from the real experimental datasets from MissionBio (https://portal.missionbio.com/datasets). We further randomized these coverage values using a discretized Gaussian distribution to mimic the non-uniform coverage distribution. After simulating the coverage value at each site of each cell, we generate reference and alternate read counts at each genomic locus for each cell using the Pólya urn model. For dropout positions, we begin with either a reference allele or an alternate allele. For a site with mutation, we start with a mutant allele and a wild-type allele in the urn. We randomly choose an allele from the urn, copy it, and return both the allele and its copy to the urn until the total coverage is reached. To introduce false positive errors at a genomic site, we perform a Pólya urn round where we return an allele different from the selected allele to the urn. To increase the likelihood of significant FP alleles being introduced into the read counts at the selected site, we sample such Pólya urn round using a linear probability mass function with a negative slope over the range [1, *coverage* + 1], ensuring *P* (*coverage* + 1) = 0. Lastly, we introduced sequencing errors caused by sequencer misinterpretations with a probability of δ = 0.001.

For modeling the copy number alterations (CNAs), we directly adopted the simulation strategy from SCIPhIN [22], where we assigned different copy numbers for different regions in the genome at the root node. We progressively lowered the probability of more copies as each additional copy is less likely. A comprehensive description of the read count simulation process is available in Supplementary Note 3.

### Evaluation metrics

To evaluate PHALCON’s performance against that of the other methods, we used four metrics, three of which evaluate the accuracy of the mutation calls, and the fourth metric evaluates the accuracy of the inferred phylogeny.

To measure variant calling accuracy, we compute recall, precision, and F1 score between the true and inferred genotype binary matrices with entries 0 (no mutation) and 1 (mutation). Precision calculates the fraction of the predicted positives that were actually positive, whereas recall finds what fraction of actual positive instances were correctly predicted. Precision and recall are computed as:

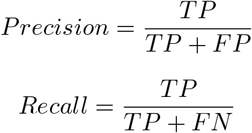

Here, TP (True Positive, actual genotype = 1, predicted genotype = 1) denotes sites that are actually mutated and also called as a mutation by the method. TN (True Negative, actual genotype = 0, predicted genotype = 0) denotes sites that do not harbor mutations and was predicted to not harbor mutation by the method. FP (False Positive, actual genotype = 0, predicted genotype = 1) indicates sites which do not harbor mutation but the method falsely predicted a mutation. FN (False Negative, actual genotype = 1, predicted genotype = 0) denotes the sites that harbor a mutation but was falsely predicted to not harbor mutation by the method.

A higher recall means that the model has a lower rate of false negatives, whereas a higher precision means that the model has a lower rate of false positives. For a complete overview of the model’s correctness, we further compute the harmonic mean of precision and recall, called F1 score, which takes into account the importance of both metrics and is calculated as:

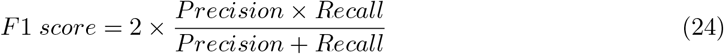

For calculating the accuracy of the inferred phylogeny, we computed MP3 similarity [28] between the inferred and ground truth phylogeny. MP3 is a triplet-based similarity that finds the number of intersecting pairs of triplets between two trees with disparate sets of mutations and is thus suitable for comparing two phylogenies whose branches are labeled with mutations. MP3 similarity also allows for poly-occurring (same label present across multiple nodes, e.g., mutation recurrence) as well as multi-labeled trees (single node having multiple labels, e.g., two different mutations occurring simultaneously). MP3 is calculated as:

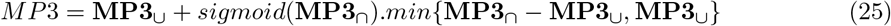

where **MP3**_∪_ is the similarity computed between all triplets of labels taken from both trees and **MP3**_∩_ is the similarity computed between the triplets of labels common between two trees. *sigmoid* is the sigmoid function, *sigmoid*(*x*) = (1 + *e*^−*ξ*(*x*−0.5)^)^−1^ centered at 0.5 with parameter *ξ* that adjusts the slopeness of the curve. Following [28], we used *ξ* = 10 in our experiments.

### Removal of germline variants from TNBC datasets

For the real tumor datasets, to ensure that germline mutations are excluded and only somatic mutations are reported, we performed post-processing steps. For the TNBC samples, we used bulk normal profiles from each patient to eliminate heterozygous germline variants. The bulk exome sequencing data from matched normal were used for filtering germline variants. The bulk normal fastq files were aligned to hg19 reference genome using Bowtie 2.0 [55] with default parameters. The resulting SAM files were converted to mpileup files using the mpileup function of SAMtools [56]. For a variant reported by PHALCON, if bulk normal mpileup harbored the variant allele (VAF threshold of 0.1) at the same site, the site was removed as a germline variant. We further utilized NCBI’s publicly available database of genetic variation, dbSNP (v138) [57] to remove known SNPs from our mutation sets. If a mutation was reported by PHALCON to be present in all cells and had evidence in dbSNP, it was removed.

### Removal of germline variants from AML datasets

For AML samples, matched bulk normal was not available. We reported the final set of variants after two post-processing steps. We first used NCBI’s publicly available database of genetic variation, dbSNP (v138) [57] to remove known SNPs from our mutation sets. If a mutation was reported by PHALCON to be present in all cells and had evidence in dbSNP, it was removed. We further removed the redundant set of intronic small insertions and deletions, particularly observed in *FLT3* gene as they appeared to be potential artifacts.

### Annotation of somatic mutations

The final set of somatic mutations reported by PHALCON were annotated using ANNOVAR [58]. ANNOVAR requires users to specify a database for gene definitions (e.g., RefSeq, UCSC Gene, etc.) and maps transcripts to the genome, establishes the relationship between transcripts and genes and finally categorizes the mutations into different classes including nonsynonymous, synonymous, stopgain, stoploss, intronic and noncoding mutations. In our analysis, we used the human genome reference build hg19 (GRCh37) to annotate the inferred SNVs and indels. After annotating the genes using ANNOVAR, we proceeded to identify if the mutations were present in known tumor suppressor genes and oncogenes. To curate a list of tumor suppressor genes and oncogenes, we utilized multiple databases: Cosmic (http://cancer.sanger.ac.uk/cosmic) database [59], the cancer gene census [60] (http://cancer.sanger.ac.uk/census), Onkogendatenbank by ZKBS BVL (https://zkbs-online.de/en/databases/oncogenes), the ONGene database (https://bioinfo-minzhao.org/ongene/) [61], and the TSGene database (https://bioinfo.uth.edu/TSGene/) [62].

### Validation of somatic mutations using bulk tumor data for TNBC

For the real tumor datasets, we used bulk tumor sequencing data from the same patient (whenever available) to validate the mutations inferred by PHALCON. For the TNBC MPT-seq datasets, we used the set of bulk tumor variants for all five patients reported by the original study (available at https://github.com/navinlabcode/MissionBio_TNBC_paper/blob/main/metadata/Total_Submitted_Genes.csv) as the reference bulk variants against which we compared the variants inferred by PHALCON.

### Validation of somatic mutations using bulk tumor data for AML

For the AML datasets, we used the publicly accessible bulk tumor samples (whichever were available) at https://www.ncbi.nlm.nih.gov/Traces/study/?acc=PRJNA648656 as the reference against which we compared the PHALCON-inferred mutations. We performed orthogonal validation analysis for SNVs and small insertions/deletions. For SNVs, we reported a mutation to be bulk-validated if its variant allele frequency (VAF) in bulk data was greater than 0.1. For insertions, we checked for the presence of the alternate sequence for it to be reported as bulk-validated. For deletion, since mpileup gives the sequence at every position where deletion has occurred, we computed the average of the VAFs across all the deleted sites (with the alternate character being “*”), and if the average VAF across those sites was greater than 0.02, we reported that deletion to be bulk-validated.

### Validation of mutations without support in tumor bulk using statistical test

For some of the mutations inferred by PHALCON from the real tumor single-cell datasets, there was no evidence in the bulk tumor data due to lack of coverage or low coverage depth. To investigate whether these inferred mutations were indeed real or caused by false positive error, we designed a statistical test. To perform this test for a site *i*, we compare two models, the inference model of PHALCON (*I*M^*P HALCON*^) and a model (*I*M^*F P*^) that assumes that the mutation at site *i* is caused by false positive error. To compare the two models, we compute the Bayes factor (BF) [21] based on the read count data at site *i*, (𝒜_*i*_, 𝒟_*i*_) given by:

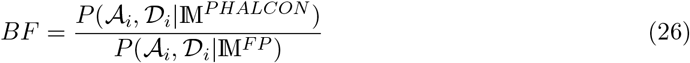

When IM^*P HALCON*^ fits the data better than *I*M^*F P*^, then *BF* becomes greater than 1 and larger value of *BF* indicates stronger evidence in favor of *I*M^*P HALCON*^ .

For IM^*F P*^, we need a likelihood function that directly models the false positive error rate using an explicit parameter. To model the false positive error rate, we adopted the genotype likelihood functions from SCIPhIN which uses a beta binomial distribution parameterized by a frequency parameter *f* and overdispersion parameter *ω* to model the read count distribution (see equations (5)-(8) in [22]). To calculate the highest possible value of *f* for a dataset (*f*_*sample*_), we selected the highest reported per-base false discovery rate in single-cell studies [10], 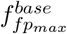 and scaled it using the total number of sites in a dataset (𝒮_*t*_) and the number of mutated sites (*n*) as given by:

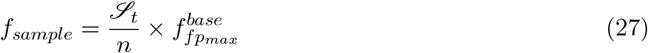

Using the *f*_*sample*_ as calculated in equation (27) and the default values of *ω* and other parameters as suggested in [22], we computed the genotype likelihoods (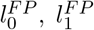,and 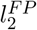 for genotypes 0, 1, 2 respectively) under the model *I*M^*F P*^ . Since *I*M^*F P*^ assumes that all the cells for site *i* are unmutated and any evidence of mutation are caused by false positive error, *P* (𝒜_*i*_, 𝒟_*i*_|*I*M^*F P*^) can be approximated as:

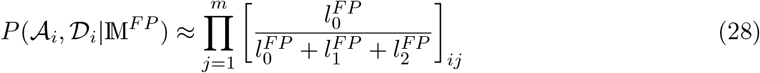

On the other hand, following [21], we approximate *P* (𝒜_*i*_, 𝒟_*i*_|*I*M^*P HALCON*^) based on the best tree (𝒯 ^*best*^) and mutation history (*σ*^*best*^) inferred by PHALCON:

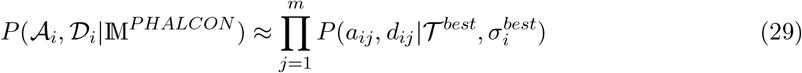

To avoid overflow, the computation of *BF* is performed in the log-space and we compute log(*BF*).

### Estimating the functional impact of mutations

To predict the functional impact of the mutations, we used CADD (Combined Annotated Dependent Depletion) [35], which employs a machine learning model to integrate 63 genomic features for prioritizing causal variants. CADD score ranks the genes in the order of increasing deleteriousness, i.e., a higher CADD score implies a more detrimental effect. We used the GRCh37-v1.7 version of the tool to obtain the CADD scores. Out of all the mutations inferred by PHALCON, we selected the top deleterious genes with highest potential functional impact on the basis of CADD scores scaled by CADD *>* 15 (top 5%).

We further utilized MutPred2 [33] for investigating the pathogenicity of specific somatic mutations. MutPred2 probabilistically scores the pathogenicity of an amino acid substitution by integrating genetic and molecular data and enables the inference of altered molecular mechanisms by assessing the impact of the mutation on more than 50 distinct protein properties. MutPred2 inferred score *>* 0.5 indicates a high pathogenicity of the mutation. For the selected mutations, we extracted the protein sequences from the UniProt database [63] in the FASTA format to provide the input to MutPred2 and MutPred2 was run with default p-value threshold of 0.05.

### Visualization of genes harboring mutations

To visualize amino acid changes in protein structures caused by mutations inferred by PHALCON, we employed the maftools (v2.18.0) [64] library in R to generate lollipop plots.

### Differential expression analysis of *PROCR* gene using TNBC scRNA-seq data

To understand the differential expression pattern of *PROCR* gene in cancer and normal epithelial cells in breast, we analyzed single-cell RNA-seq data from TNBC patients from a single-cell atlas of breast cancer [65]. 100064 single-cells from 10 TNBC patients were integrated using scDREAMER [66] to infer the latent embeddings of the cells by eliminating the batch effect. The epithelial cells were selected for further analysis and the expression of *PROCR* gene was compared across the cancer and normal epithelial cells using t-test and visualized using the ‘pl.violin()’ function of Scanpy [67]. Scanpy’s UMAP function was used for plotting the cellular embeddings in 2D.

### Survival analysis for new mutations detected by PHALCON in TNBC and AML cohorts

We performed a survival analysis for the *PROCR* gene using clinical and expression data of 89 TNBC patients from the TCGA (The Cancer Genome Atlas) cohort [34]. Patients were classified into two groups (high and low) based on the median value of *PROCR* expression. For survival analysis of genes for which PHALCON reported more mutations in the AML cohort, we integrated the clinical and mutational data of two large AML cohorts. The first cohort, the Oregon Health & Science University (OHSU) Beat AML 2.0 cohort [37], consisted of 942 samples from 805 patients, and the second cohort, the TCGA AML cohort [38], consisted of 200 samples from 200 patients, resulting in a cumulative cohort of 1005 patients. Patients were classified into two groups (mutant and wildtype) based on the presence of mutation in the target gene. Survival data was generated using the R “survival” package [68] and visualized as Kaplan-Meier plots using the R “survminer” package [69]. We used log-rank tests to calculate the p-values.

### Derivation of pathway mutation profiles for AML samples

To generate the pathway mutation profile of an AML patient in the 123 AML patients cohort, we first selected the top 20 mutated genes detected by PHALCON. These 20 genes were associated with seven major pathways including DNA methylation, receptor tyrosine kinase (RTK)/Ras GTPase (RAS)/MAP Kinase (MAPK) signaling (RTK-RAS), chromatin/cohesin, *NPM1*, transcription factor, splicing and apoptosis (Supplementary Table 2). For determining the pathway mutation profile for each patient, the genes from one pathway were grouped together and their cellular mutation profiles were merged into a single mutation profile for the pathway where an entry was set to 1 if at least one of the genes in the pathway was mutated otherwise it was set to 0 if none of the pathway genes were mutated. Using this single mutation profile, the fraction of cells mutated for the specific pathway was calculated. Finally, the pathway mutation profile for a patient sample was a 7-dimensional vector where each dimension represented the fraction of cells mutated for a specific pathway.

### Mapping of patient samples in the combined AML cohort to pathway subgroups defined based on 123 AML samples

Once we obtain the pathway mutation profiles for each patient in the smaller cohort of 123 patients, we utilized the pathway mutation profiles to cluster the samples into pathway subgroups. The clustering was performed using the Leiden algorithm by employing the “sc.tl.leiden” function of Scanpy [67] which identified 5 pathway subgroups, each characterized uniquely by a combination of the fraction of cells mutated in each of the 7 pathways. We then mapped the patient samples in the combined cohort of AML curated using the TCGA cohort consisting of 200 samples from 200 patients [38], and the BeatAML 2.0 cohort consisting of 942 samples from 805 patients [37] to the 5 pathway subgroups. To do so, first the pathway mutation profile for each sample in the combined cohort was calculated by computing the mean VAF across genes corresponding to the respective pathways. After that, for each sample, we computed the Euclidean distance of its pathway mutation profile to the centroids of the five pathway subgroups. The sample was assigned to the pathway subgroup for which distance was minimum. After assignment of all the patient samples in the combined cohort, we identified 5 pathway subgroups.

### Survival analysis of pathway subgroups in the combined AML cohort

Using the pathway subgroups obtained from the combined AML cohort of 1005 patients, we performed overall survival analysis, group-wise (one subgroup vs others) survival analysis and subgroup to subgroup (i.e., one subgroup vs another subgroup) survival analysis. Survival data was generated using the R “survival” package [68] and visualized as Kaplan-Meier plots using the R “survminer” package [69]. We used log-rank tests to calculate the p-values.

### Calculation of log odds ratio for determining cell-level co-occurrence and mutual exclusivity

To evaluate the pairwise association between different mutations inferred by PHALCON for a sample, we first computed a contingency table using the genotype matrix to find the number of cells with mutant or wild-type genes. Based on the contingency table, for each pair of genes, the log odds ratio was calculated. We corrected the ratio using Haldane correction (δ = 0.5). We then evaluated the significance of the association by calculating p-value on the contingency table using Fisher’s exact test. Fisher’s exact test was performed using the”fisher.test” function with default settings in R.

## Data Availability

The TNBC tumor datasets (bulk normal, bulk tumor, and single-cell sequencing) are publicly available at the NCBI Sequencing Read Archive (SRA) under the accession SRA: PRJNA763862 (https://www.ncbi.nlm.nih.gov/sra/?term=PRJNA763862). The data from TCGA BRCA cohort used for survival analysis are publicly available under the dbGAP study accession phs000178 (https://portal.gdc.cancer.gov/projects/TCGA-BRCA). The processed scRNA-seq data from the breast cancer patients are available through the Broad Institute Single Cell portal at https://singlecell.broadinstitute.org/single_cell/study/SCP1039 as well as through the Gene Expression Omnibus under accession number GSE176078. The AML tumor datasets (bulk tumor data, single-cell sequencing data) are publicly available at the NCBI BioProject ID under the accession SRA: PRJNA648656 (https://www.ncbi.nlm.nih.gov/Traces/study/?acc=PRJNA648656). Clinical and mutation data of the patients from the BeatAML 2.0 cohort are available at https://biodev.github.io/BeatAML2/. Clinical and mutation data of the patients from the TCGA AML cohort are publicly available at https://gdc.cancer.gov/about-data/publications/laml_2012.

## Code Availability

PHALCON is implemented in Python 3 and is freely available at https://github.com/Zafar-Lab/PHALCON under the MIT license.

## Acknowledgements

We thank Prof. Nicholas Navin and Jake Leighton from MD Anderson Cancer Center for constructive discussions and for providing us with the loom files for the TNBC cohort. We also thank Prof.

Koichi Takahashi from MD Anderson Cancer Center for kindly providing us with the loom files for the AML cohort.

## Author contributions

H.Z. designed the study. P. and H.Z. developed the model and algorithm. P. and S.C. implemented the method. P. conducted all the experiments. P. and A.G. conducted the survival analyses for real datasets. P. and H.Z. wrote the manuscript and all authors approved the manuscript.

## References

[1] Navin, N. et al. Tumour evolution inferred by single-cell sequencing. Nature 472, 90–94 (2011). URL 10.1038/nature09807.

[2] Kashima, Y. et al. Single-cell sequencing techniques from individual to multiomics analyses. Experimental & Molecular Medicine 52, 1419–1427 (2020). URL 10.1038/s12276-020-00499-2.

[3] Lim, B., Lin, Y. & Navin, N. Advancing cancer research and medicine with single-cell genomics. Cancer Cell 37, 456–470 (2020). URL https://www.sciencedirect.com/science/article/pii/S1535610820301483.

[4] Navin, N. E. Cancer genomics: one cell at a time. Genome Biology 15, 452 (2014). URL 10.1186/s13059-014-0452-9.

[5] Cui, Y. et al. Single-cell transcriptome analysis maps the developmental track of the human heart. Cell Reports 26, 1934–1950.e5 (2019). URL https://www.sciencedirect.com/science/article/pii/S2211124719301081.

[6] Navin, N. E. The first five years of single-cell cancer genomics and beyond. Genome Research 25, 1499–1507 (2015). URL http://genome.cshlp.org/content/25/10/1499.abstract. http://genome.cshlp.org/content/25/10/1499.full.pdf+html.

[7] Burrell, R. A., McGranahan, N., Bartek, J. & Swanton, C. The causes and consequences of genetic heterogeneity in cancer evolution. Nature 501, 338–345 (2013). URL 10.1038/nature12625.

[8] Wu, X. et al. Clonal selection drives genetic divergence of metastatic medulloblastoma. Nature 482, 529 EP – (2012). URL 10.1038/nature10825.

[9] Merlo, L. M., Pepper, J. W., Reid, B. J. & Maley, C. C. Cancer as an evolutionary and ecological process. Nature Reviews Cancer 6, 924–935 (2006). URL 10.1038/nrc2013.

[10] Zafar, H., Navin, N., Nakhleh, L. & Chen, K. Computational approaches for inferring tumor evolution from single-cell genomic data. Current Opinion in Systems Biology 7, 16–25 (2018). URL https://www.sciencedirect.com/science/article/pii/S2452310017301865. † Future of systems biology† Genomics and epigenomics.

[11] McKenna, A. et al. The Genome Analysis Toolkit: A MapReduce framework for analyzing next-generation DNA sequencing data. Genome Research 20, 1297–1303 (2010). URL http://genome.cshlp.org/content/20/9/1297.abstract. http://genome.cshlp.org/content/20/9/1297.full.pdf+html.

[12] Zafar, H., Wang, Y., Nakhleh, L., Navin, N. & Chen, K. Monovar: single-nucleotide variant detection in single cells. Nature Methods 13, 505–507 (2016). URL 10.1038/nmeth.3835.

[13] Dong, X. et al. Accurate identification of single-nucleotide variants in whole-genome-amplified single cells. Nature Methods 14, 491–493 (2017). URL 10.1038/nmeth.4227.

[14] Jahn, K., Kuipers, J. & Beerenwinkel, N. Tree inference for single-cell data. Genome Biology 17, 86 (2016). URL 10.1186/s13059-016-0936-x.

[15] Zafar, H., Tzen, A., Navin, N., Chen, K. & Nakhleh, L. SiFit: inferring tumor trees from single-cell sequencing data under finite-sites models. Genome Biology 18, 178 (2017). URL 10.1186/s13059-017-1311-2.

[16] Zafar, H., Navin, N., Chen, K. & Nakhleh, L. SiCloneFit: Bayesian inference of population structure, genotype, and phylogeny of tumor clones from single-cell genome sequencing data. Genome Research 29, 1847–1859 (2019). URL http://genome.cshlp.org/content/29/11/1847.abstract. http://genome.cshlp.org/content/29/11/1847.full.pdf+html.

[17] El-Kebir, M. SPhyR: tumor phylogeny estimation from single-cell sequencing data under loss and error. Bioinformatics 34, i671–i679 (2018). URL 10.1093/bioinformatics/bty589.

[18] Singer, J., Kuipers, J., Jahn, K. & Beerenwinkel, N. Single-cell mutation identification via phylogenetic inference. Nature Communications 9, 5144 (2018). URL 10.1038/s41467-018-07627-7.

[19] Edrisi, M., Zafar, H. & Nakhleh, L. A Combinatorial Approach for Single-cell Variant Detection via Phylogenetic Inference. In Huber, K.T. & Gusfield, D. (eds.) 19th International Workshop on Algorithms in Bioinformatics (WABI 2019), vol. 143 of Leibniz International Proceedings in Informatics (LIPIcs), 22:1–22:13 (Schloss Dagstuhl–Leibniz-Zentrum fuer Informatik, Dagstuhl, Germany, 2019). URL http://drops.dagstuhl.de/opus/volltexte/2019/11052.

[20] Edrisi, M. et al. Phylovar: toward scalable phylogeny-aware inference of single-nucleotide variations from single-cell DNA sequencing data. Bioinformatics 38, i195–i202 (2022). URL 10.1093/bioinformatics/btac254.

[21] Kuipers, J., Jahn, K., Raphael, B. J. & Beerenwinkel, N. Single-cell sequencing data reveal widespread recurrence and loss of mutational hits in the life histories of tumors. Genome Research 27, 1885–1894 (2017). URL http://genome.cshlp.org/content/27/11/1885.abstract. http://genome.cshlp.org/content/27/11/1885.full.pdf+html.x

[22] Kuipers, J., Singer, J. & Beerenwinkel, N. Single-cell mutation calling and phylogenetic tree reconstruction with loss and recurrence. Bioinformatics 38, 4713–4719 (2022). URL 10.1093/bioinformatics/btac577.

[23] Kang, S. et al. SIEVE: joint inference of single-nucleotide variants and cell phylogeny from single-cell DNA sequencing data. Genome Biology 23, 248 (2022). URL 10.1186/s13059-022-02813-9.

[24] Morita, K. et al. Clonal evolution of acute myeloid leukemia revealed by high-throughput single-cell genomics. Nature Communications 11, 5327 (2020). URL 10.1038/s41467-020-19119-8.

[25] Miles, L. A. et al. Single-cell mutation analysis of clonal evolution in myeloid malignancies. Nature 587, 477–482 (2020). URL 10.1038/s41586-020-2864-x.

[26] Yaeger, R. et al. Molecular Characterization of Acquired Resistance to KRASG12C–EGFR Inhibition in Colorectal Cancer. Cancer Discovery 13, 41–55 (2023). URL 10.1158/2159-8290.CD-22-0405. https://aacrjournals.org/cancerdiscovery/article-pdf/13/1/41/3351061/41.pdf.

[27] Leighton, J., Hu, M., Sei, E., Meric-Bernstam, F. & Navin, N. E. Reconstructing mutational lineages in breast cancer by multi-patient-targeted single-cell DNA sequencing. Cell Genomics 3, 100215 (2023). URL https://www.sciencedirect.com/science/article/pii/S2666979X22001689.

[28] Ciccolella, S. et al. Triplet-based similarity score for fully multilabeled trees with poly-occurring labels. Bioinformatics 37, 178–184 (2021). URL 10.1093/bioinformatics/btaa676.

[29] Sollier, E., Kuipers, J., Takahashi, K., Beerenwinkel, N. & Jahn, K. COMPASS: joint copy number and mutation phylogeny reconstruction from amplicon single-cell sequencing data. Nature Communications 14, 4921 (2023). URL 10.1038/s41467-023-40378-8.

[30] Wong, K.-B. et al. Hot-spot mutants of p53 core domain evince characteristic local structural changes. Proceedings of the National Academy of Sciences 96, 8438–8442 (1999). URL 10.1073/pnas.96.15.8438.

[31] Feng, M. et al. RASAL2 activates RAC1 to promote triple-negative breast cancer progression. The Journal of Clinical Investigation 124, 5291–5304 (2014). URL https://www.jci.org/articles/view/76711.

[32] Wang, D. et al. Protein C receptor is a therapeutic stem cell target in a distinct group of breast cancers. Cell Research 29, 832–845 (2019). URL 10.1038/s41422-019-0225-9.

[33] Pejaver, V. et al. Inferring the molecular and phenotypic impact of amino acid variants with MutPred2. Nature Communications 11, 5918 (2020). URL 10.1038/s41467-020-19669-x.

[34] Ciriello, G. et al. Comprehensive Molecular Portraits of Invasive Lobular Breast Cancer. Cell 163, 506–519 (2015). URL https://www.sciencedirect.com/science/article/pii/S0092867415011952.

[35] Rentzsch, P., Witten, D., Cooper, G. M., Shendure, J. & Kircher, M. CADD: predicting the deleteriousness of variants throughout the human genome. Nucleic Acids Research 47, D886–D894 (2018). URL 10.1093/nar/gky1016. https://academic.oup.com/nar/article-pdf/47/D1/D886/27436395/gky1016.pdf.

[36] Sportoletti, P., Sorcini, D. & Falini, B. Bcor gene alterations in hematologic diseases. Blood 138, 2455–2468 (2021). URL 10.1182/blood.2021010958. https://ashpublications.org/blood/article-pdf/138/24/2455/1853492/bloodbld2021010958.pdf.

[37] Bottomly, D. et al. Integrative analysis of drug response and clinical outcome in acute myeloid leukemia. Cancer Cell 40, 850–864.e9 (2022). URL https://www.sciencedirect.com/science/article/pii/S1535610822003129.

[38] Network, T. C. G. A. R. Genomic and epigenomic landscapes of adult de novo acute myeloid leukemia. New England Journal of Medicine 368, 2059–2074 (2013). URL 10.1056/NEJMoa1301689.

[39] Schwede, M. et al. Mutation order in acute myeloid leukemia identifies uncommon patterns of evolution and illuminates phenotypic heterogeneity. Leukemia 38, 1501–1510 (2024).

[40] Döhner, H. et al. Diagnosis and management of aml in adults: 2022 recommendations from an international expert panel on behalf of the eln. Blood 140, 1345–1377 (2022). URL 10.1182/blood.2022016867. https://ashpublications.org/blood/article-pdf/140/12/1345/1921355/bloodbld2022016867.pdf.

[41] Papaemmanuil, E. et al. Genomic classification and prognosis in acute myeloid leukemia. New England Journal of Medicine 374, 2209–2221 (2016). URL https://www.nejm.org/doi/full/10.1056/NEJMoa1516192. https://www.nejm.org/doi/pdf/10.1056/NEJMoa1516192.

[42] Yoshida, K. et al. Frequent pathway mutations of splicing machinery in myelodysplasia. Nature 478, 64–69 (2011). URL 10.1038/nature10496.

[43] Abelson, S. et al. Prediction of acute myeloid leukaemia risk in healthy individuals. Nature 559, 400–404 (2018). URL 10.1038/s41586-018-0317-6.

[44] Van Vlierberghe, P. et al. Phf6 mutations in adult acute myeloid leukemia. Leukemia 25, 130–134 (2011). URL 10.1038/leu.2010.247.

[45] Sportoletti, P., Sorcini, D. & Falini, B. Bcor gene alterations in hematologic diseases. Blood 138, 2455–2468 (2021). URL 10.1182/blood.2021010958.

[46] Eisfeld, A.-K. et al. Nf1 mutations are recurrent in adult acute myeloid leukemia and confer poor outcome. Leukemia 32, 2536–2545 (2018). URL 10.1038/s41375-018-0147-4.

[47] Figueroa, M. E. et al. Dna methylation signatures identify biologically distinct subtypes in acute myeloid leukemia. Cancer Cell 17, 13–27 (2010). URL https://www.sciencedirect.com/science/article/pii/S1535610809004206.

[48] Gaidzik, V. I. et al. Runx1 mutations in acute myeloid leukemia are associated with distinct clinico-pathologic and genetic features. Leukemia 30, 2160–2168 (2016). URL 10.1038/leu.2016.126.

[49] Ochi, Y. et al. Combined cohesin–runx1 deficiency synergistically perturbs chromatin looping and causes myelodysplastic syndromes. Cancer Discovery 10, 836–853 (2020). URL 10.1158/2159-8290.CD-19-0982. https://aacrjournals.org/cancerdiscovery/article-pdf/10/6/836/1814995/836.pdf.

[50] Lodato, M. A. et al. Somatic mutation in single human neurons tracks developmental and transcriptional history. Science 350, 94–98 (2015). URL 10.1126/science.aab1785.

[51] Traag, V. A., Waltman, L. & van Eck, N. J. From Louvain to Leiden: guaranteeing well-connected communities. Scientific Reports 9, 5233 (2019). URL 10.1038/s41598-019-41695-z.

[52] Saitou, N. & Nei, M. The neighbor-joining method: a new method for reconstructing phylogenetic trees. Molecular Biology and Evolution 4, 406–425 (1987). URL 10.1093/oxfordjournals.molbev.a040454.

[53] Mäkinen, E. & Siltaneva, J. A Note on Rémy’s Algorithm for Generating Random Binary Trees. Missouri Journal of Mathematical Sciences 15, 103–109 (2003).

[54] Salehi, S. et al. ddClone: joint statistical inference of clonal populations from single cell and bulk tumour sequencing data. Genome Biology 18, 44 (2017). URL 10.1186/s13059-017-1169-3.

[55] Langmead, B. & Salzberg, S. L. Fast gapped-read alignment with Bowtie 2. Nature Methods 9, 357–359 (2012). URL 10.1038/nmeth.1923.

[56] Li, H. et al. The Sequence Alignment/Map format and SAMtools. Bioinformatics 25, 2078–2079 (2009). URL 10.1093/bioinformatics/btp352. https://academic.oup.com/bioinformatics/article-pdf/25/16/2078/48994296/bioinformatics_25_16_2078.pdf.

[57] Sherry, S. T. et al. dbSNP: the NCBI database of genetic variation. Nucleic Acids Research 29, 308–311 (2001). URL 10.1093/nar/29.1.308.

[58] Wang, K., Li, M. & Hakonarson, H. ANNOVAR: functional annotation of genetic variants from high-throughput sequencing data. Nucleic Acids Research 38, e164–e164 (2010). URL 10.1093/nar/gkq603.

[59] Forbes, S. A. et al. COSMIC: exploring the world’s knowledge of somatic mutations in human cancer. Nucleic Acids Research 43, D805–D811 (2015). URL 10.1093/nar/gku1075.

[60] Futreal, P. A. et al. A census of human cancer genes. Nature Reviews Cancer 4, 177–183 (2004). URL 10.1038/nrc1299.

[61] Liu, Y., Sun, J. & Zhao, M. ONGene: A literature-based database for human oncogenes. Journal of Genetics and Genomics 44, 119–121 (2017). URL https://www.sciencedirect.com/science/article/pii/S1673852716302053.

[62] Zhao, M., Sun, J. & Zhao, Z. TSGene: a web resource for tumor suppressor genes. Nucleic Acids Research 41, D970–D976 (2013). URL 10.1093/nar/gks937.

[63] Consortium, T. U. UniProt: the Universal Protein Knowledgebase in 2023. Nucleic Acids Research 51, D523– D531 (2022). URL 10.1093/nar/gkac1052. https://academic.oup.com/nar/article-pdf/51/D1/D523/48441158/gkac1052.pdf.

[64] Mayakonda, A., Lin, D.-C., Assenov, Y., Plass, C. & Koeffler, H. P. Maftools: efficient and comprehensive analysis of somatic variants in cancer. Genome Research 28, 1747–1756 (2018). URL http://genome.cshlp.org/content/28/11/1747.abstract. http://genome.cshlp.org/content/28/11/1747.full.pdf+html.

[65] Wu, S. Z. et al. A single-cell and spatially resolved atlas of human breast cancers. Nature Genetics 53, 1334–1347 (2021). URL 10.1038/s41588-021-00911-1.

[66] Shree, A., Pavan, M. K. & Zafar, H. scDREAMER for atlas-level integration of single-cell datasets using deep generative model paired with adversarial classifier. Nature Communications 14, 7781 (2023). URL 10.1038/s41467-023-43590-8.

[67] Wolf, F. A., Angerer, P. & Theis, F. J. SCANPY: large-scale single-cell gene expression data analysis. Genome Biology 19, 15 (2018). URL 10.1186/s13059-017-1382-0.

[68] Therneau, T. M. A Package for Survival Analysis in R (2024). URL https://CRAN.R-project.org/package=survival. R package version 3.8-3.

[69] Kassambara, A., Kosinski, M. & Biecek, P. survminer: Drawing Survival Curves using ‘ggplot2’ (2024). URL https://github.com/kassambara/survminer. R package version 0.5.0.999.

