## Supplementary Material for "PHALCON: phylogeny-aware variant calling from large-scale single-cell panel sequencing datasets"

March 4, 2025

#### Contents

**Supplementary Figures** **2**

**Supplementary Tables** **20**

**Supplementary Note 1: Description of filters used by PHALCON for identifying candidate mutation sites** **21**

**Supplementary Note 2: Description of possible mutation histories due to heterozygous, parallel, and back mutations for a given phylogeny** **23**

**Supplementary Note 3: Description of read count simulation** **25**

### Supplementary Figures

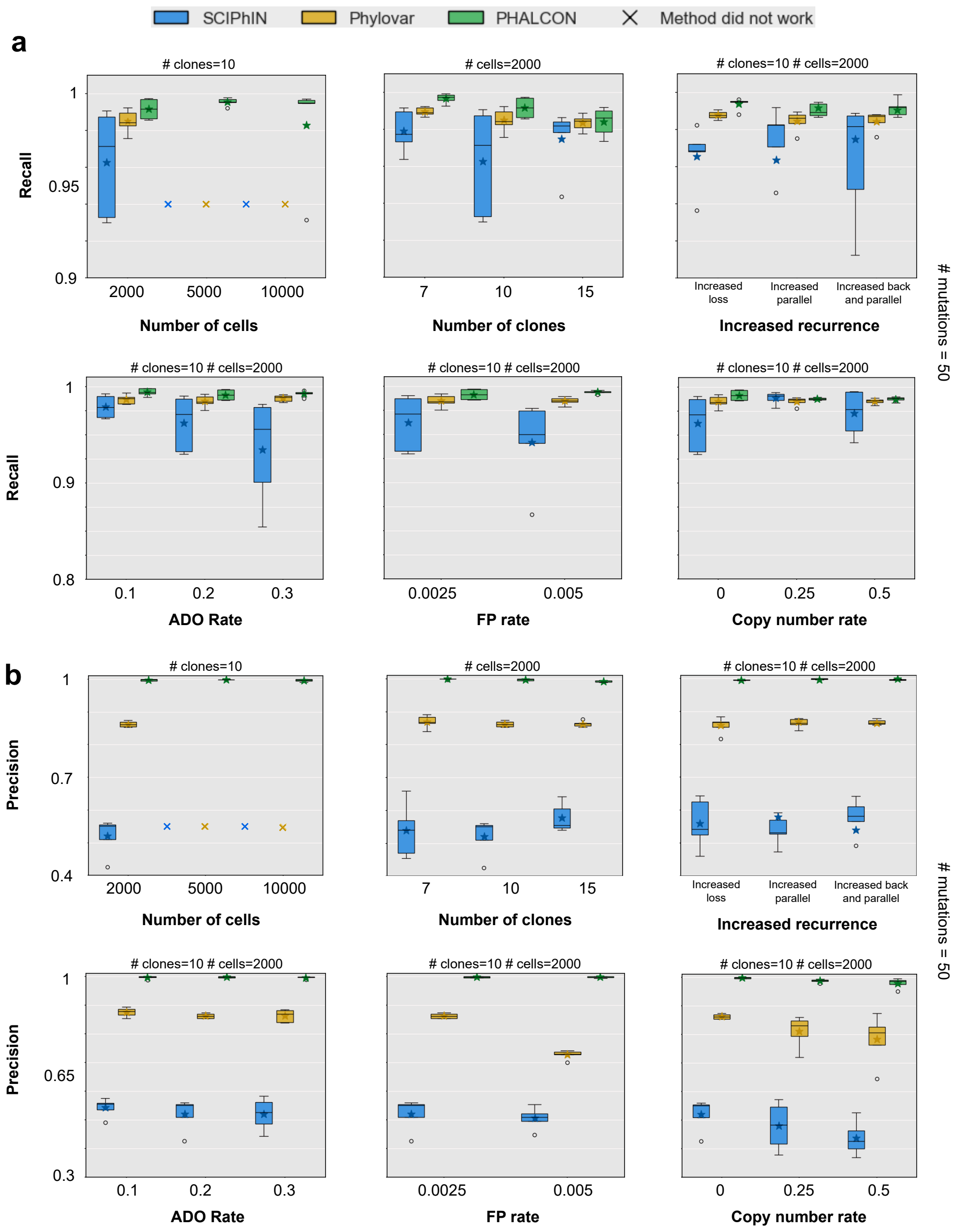


Supplementary Figure 1: Performance comparison between SCIPhIN, Phylovar and PHALCON across various experimental settings for evaluating accuracy of mutation calling. (a) Comparison in terms of recall. (b) Comparison in terms of precision. Experimental settings include varying number of cells, varying number of clones, varying rates of mutation loss and recurrence, varying ADO and FP rates, and copy number alteration rate. The simulated datasets contained 50 mutations across all experiments. A × indicates an instance where the method did not produce any result within the stipulated time (72 hours).

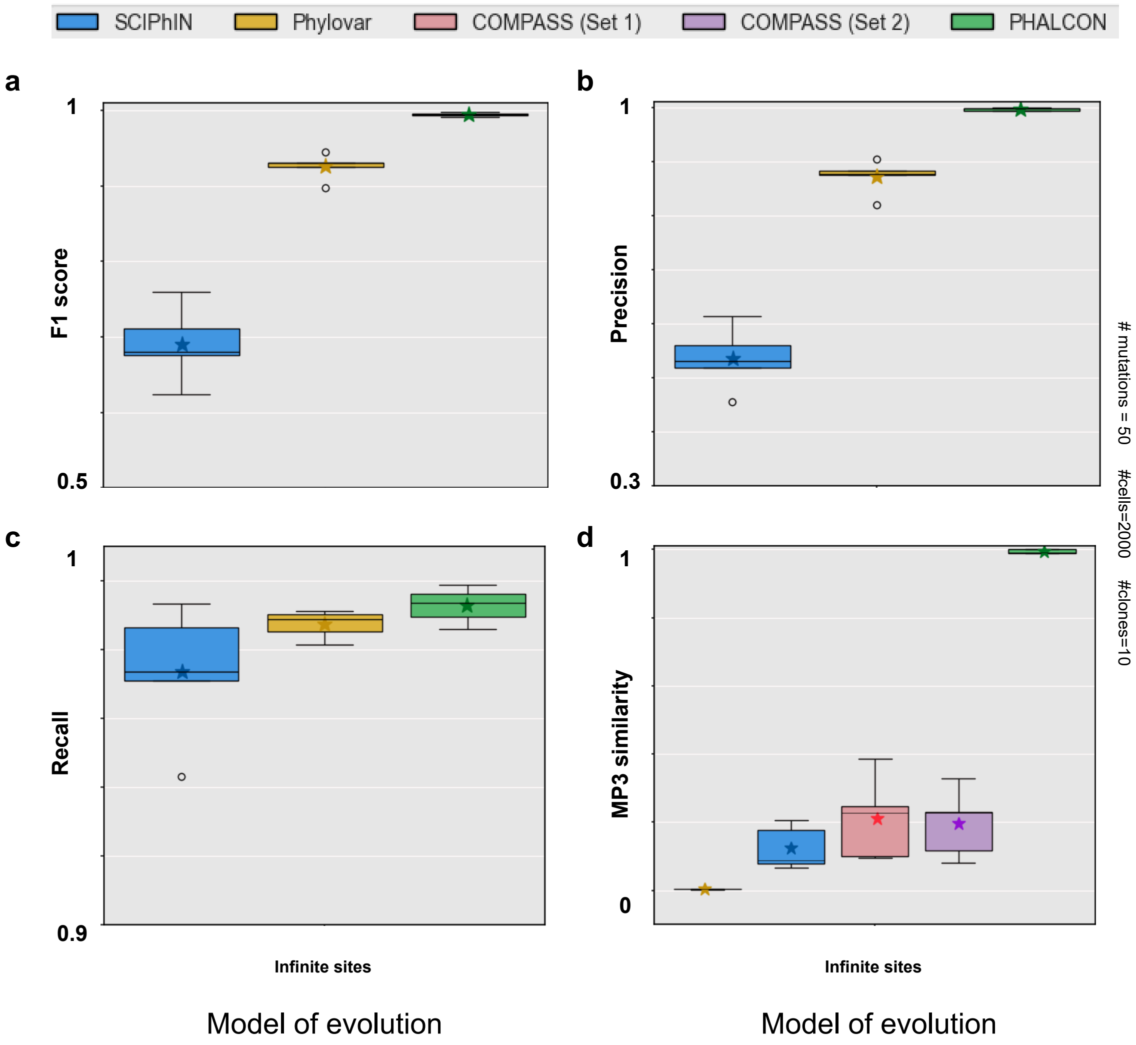


Supplementary Figure 2: Performance comparison between SCIPhIN, Phylovar, COMPASS, and PHALCON on simulated datasets generated under an infinite sites model of evolution. Comparison in terms of (a) F1 score, (b) Precision, (c) Recall, and (d) MP3 similarity. These datasets contained 2000 cells, 10 clones and 50 mutations. MP3 similarity is calculated for COMPASS (Set 1) by passing the true mutation sites as input to COMPASS for phylogeny reconstruction. For COMPASS (Set 2), only those sites are passed that are selected by COMPASS in Set 1 for reconstructing the phylogeny.

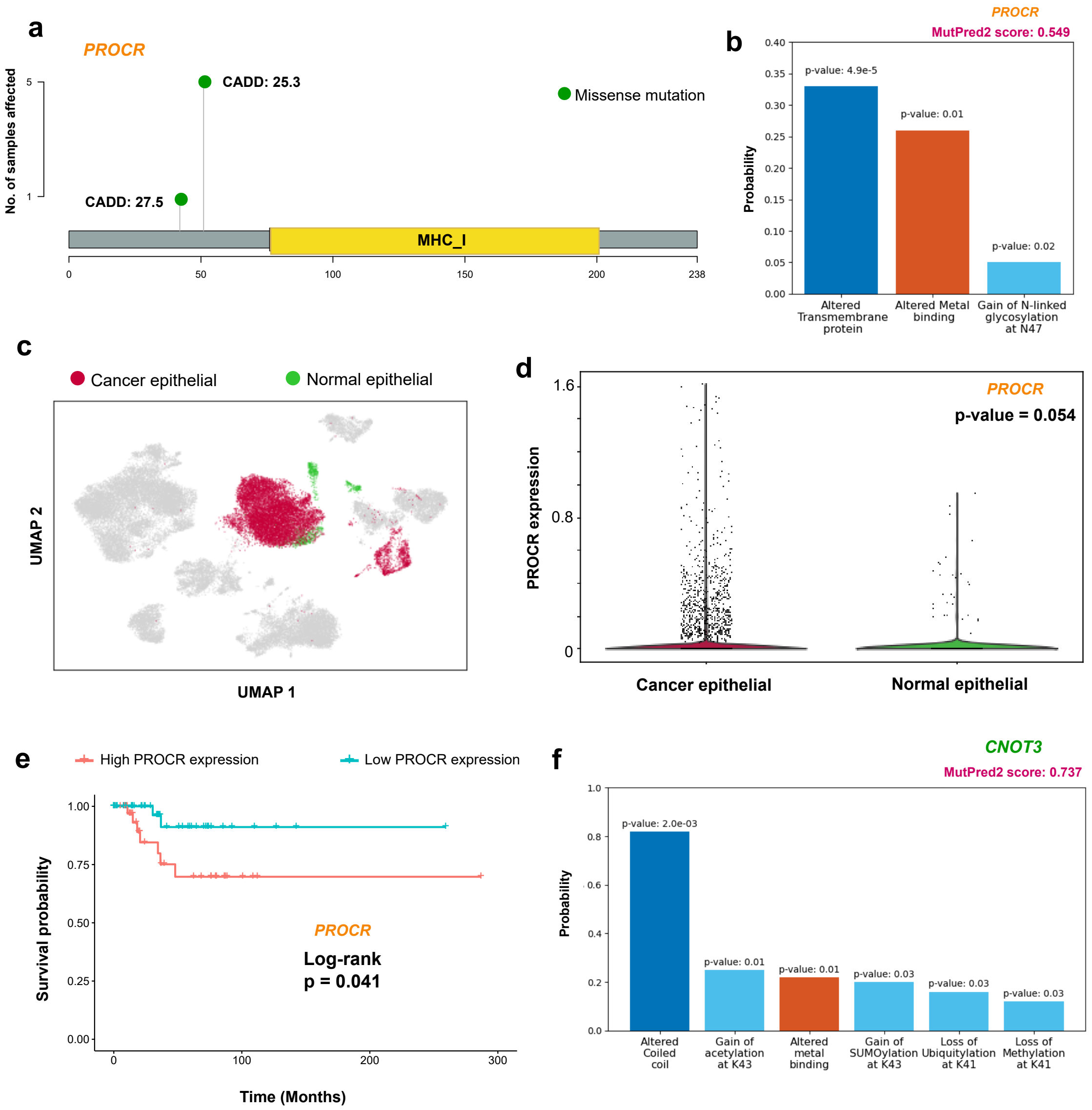


Supplementary Figure 3: (a) Lollipop plot for *PROCR* showing the amino acid changes for the two mutations (p.V42M, p.G51E) inferred by PHALCON in the TNBC cohort of 5 patients. (b) MutPred2 prediction (score = 0.549) for significantly altered molecular mechanisms (p-value $< 0.05$) for *PROCR* mutation. The bars are colored based on the ontology of the molecular mechanism. (c)-(d) Comparison of *PROCR* expression using scRNA-seq data from 10 TNBC patients from a breast cancer atlas. (c) UMAP projection and visualization of the cells based on the embeddings inferred by scDREAMER. Cancer epithelial and normal epithelial cells are highlighted in the UMAP. (d) Comparison of *PROCR* expression in cancer and normal epithelial cells. (e) Survival analysis of *PROCR* expression utilizing TCGA cohort of 89 TNBC patients. Higher *PROCR* expression is linked to poor survival with a significant p-value of 0.041. (f) MutPred2 prediction (score = 0.737) for significantly altered molecular mechanisms (p-value $< 0.05$) for *CNOT3* mutation. The bars are colored based on the ontology of the molecular mechanism.

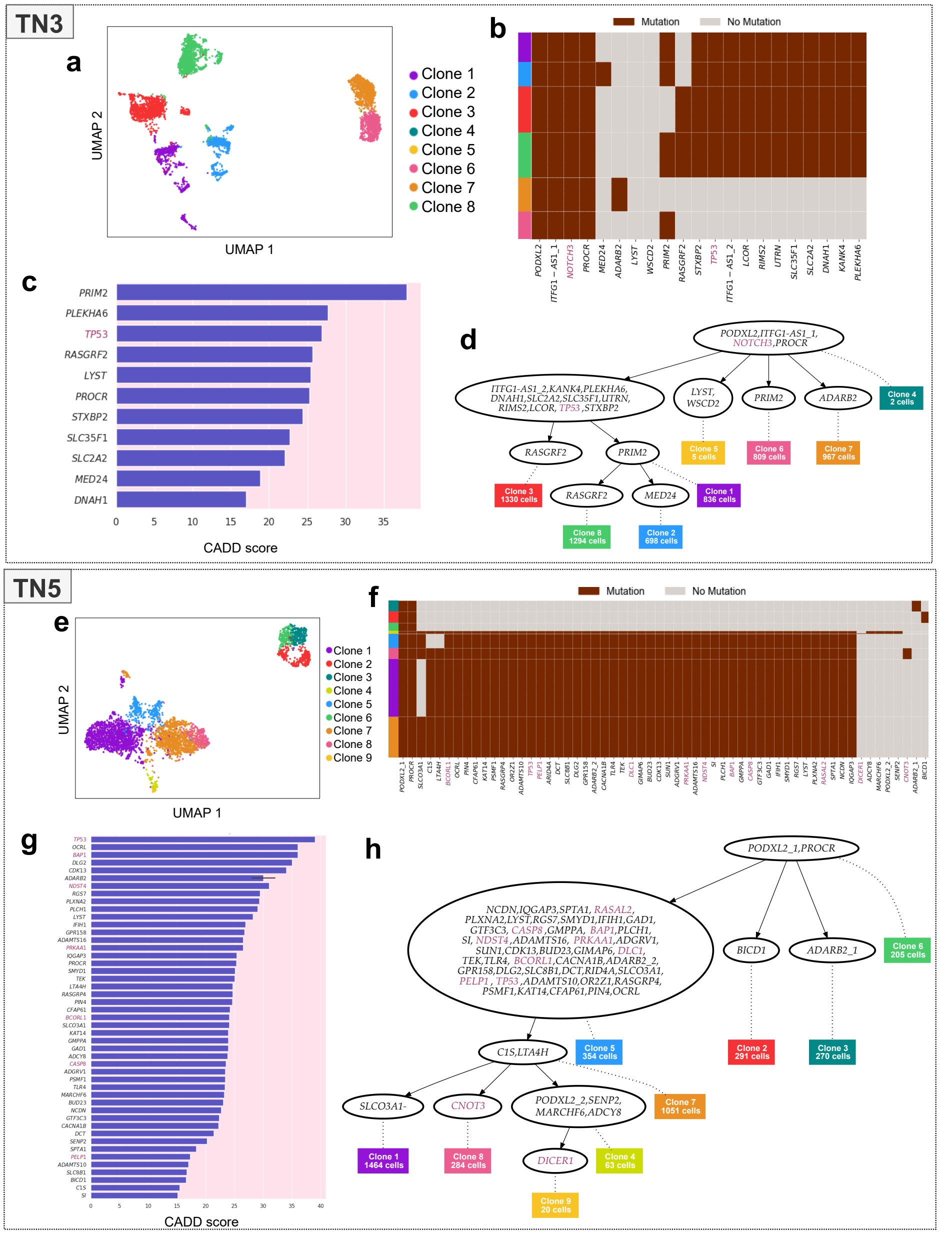


Supplementary Figure 4: Visualization of the clones inferred by PHALCON in UMAP for samples (a) TN3 and (e) TN5, respectively. (b,f) Heatmaps of clonally clustered mutations in (b) TN3 and (f) TN5, respectively as inferred by PHALCON, illustrating the clonal substructure and heterogeneity within tumor subpopulations. The clonal membership of individual cells is represented in the leftmost column. Mutations ranked according to their CADD scores denoting the functional impact of mutations' deleteriousness in samples (c) TN3 and (g) TN5 (CADD> 15, top 5%) respectively. Clonal phylogeny reconstructed by PHALCON delineating the evolutionary history of the samples (d) TN3 and (h) TN5. The leaves (rectangular boxes colored according to clone identity) represent different sub-clones. Intermediate nodes contain sub-clonal mutations that are passed down the phylogeny. The root node contains the clonal mutations present across all cells. Oncogenes and Tumor suppressor genes are highlighted in purple across all panels.

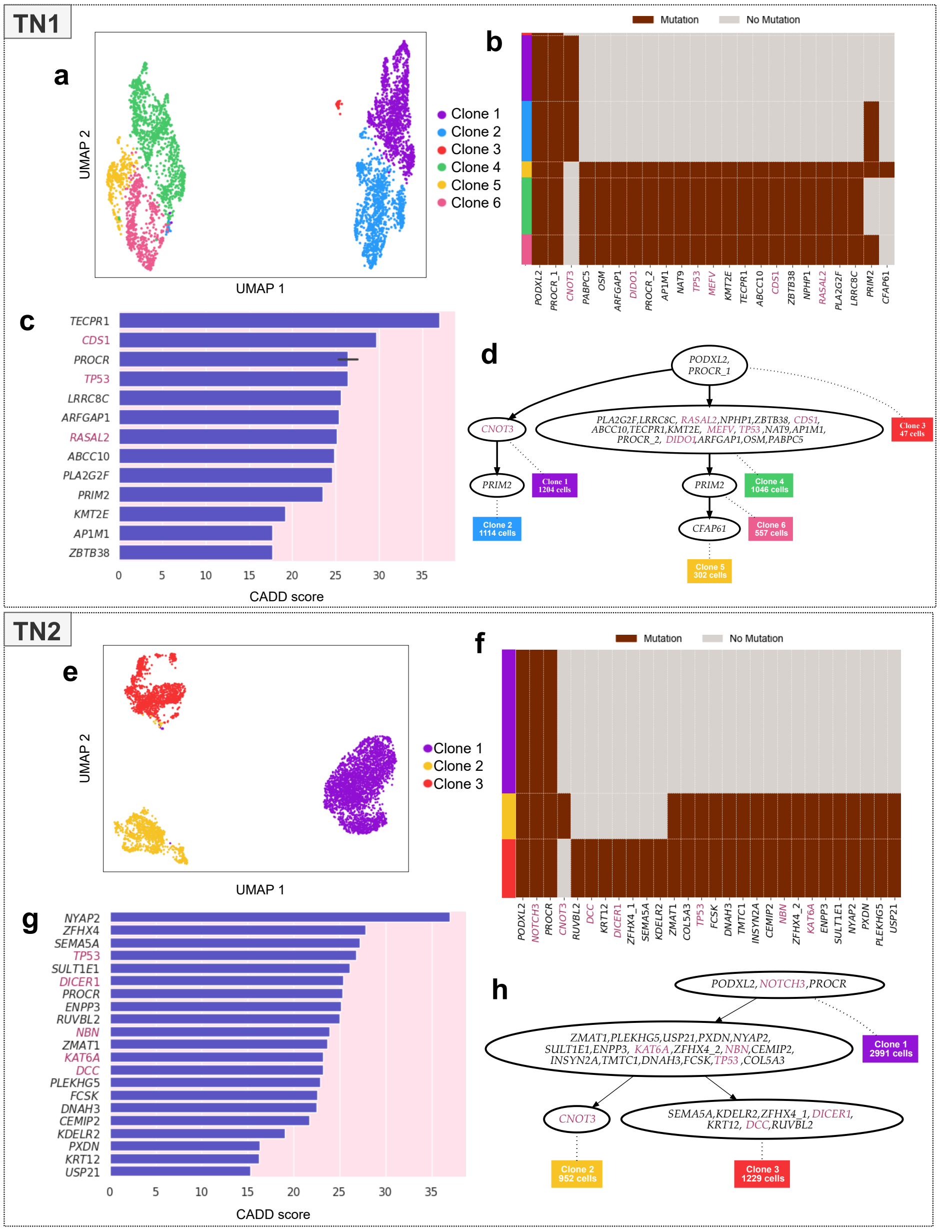

Supplementary Figure 5: Visualization of the clones inferred by PHALCON in UMAP for samples (a) TN1 and (e) TN2, respectively. (b,f) Heatmaps of clonally clustered mutations in (b) TN1 and (f) TN2, respectively as inferred by PHALCON, illustrating the clonal substructure and heterogeneity within tumor subpopulations. The clonal membership of individual cells is represented in the leftmost column. Mutations ranked according to their CADD scores denoting the functional impact of mutations' deleteriousness in samples (c) TN1 and (g) TN2 (CADD> 15, top 5%) respectively. Clonal phylogeny reconstructed by PHALCON delineating the evolutionary history of the samples (d) TN1 and (h) TN2. The leaves (rectangular boxes colored according to clone identity) represent different sub-clones. Intermediate nodes contain sub-clonal mutations that are passed down the phylogeny. The root node contains the clonal mutations present across all cells. Oncogenes and Tumor suppressor genes are highlighted in purple across all panels.

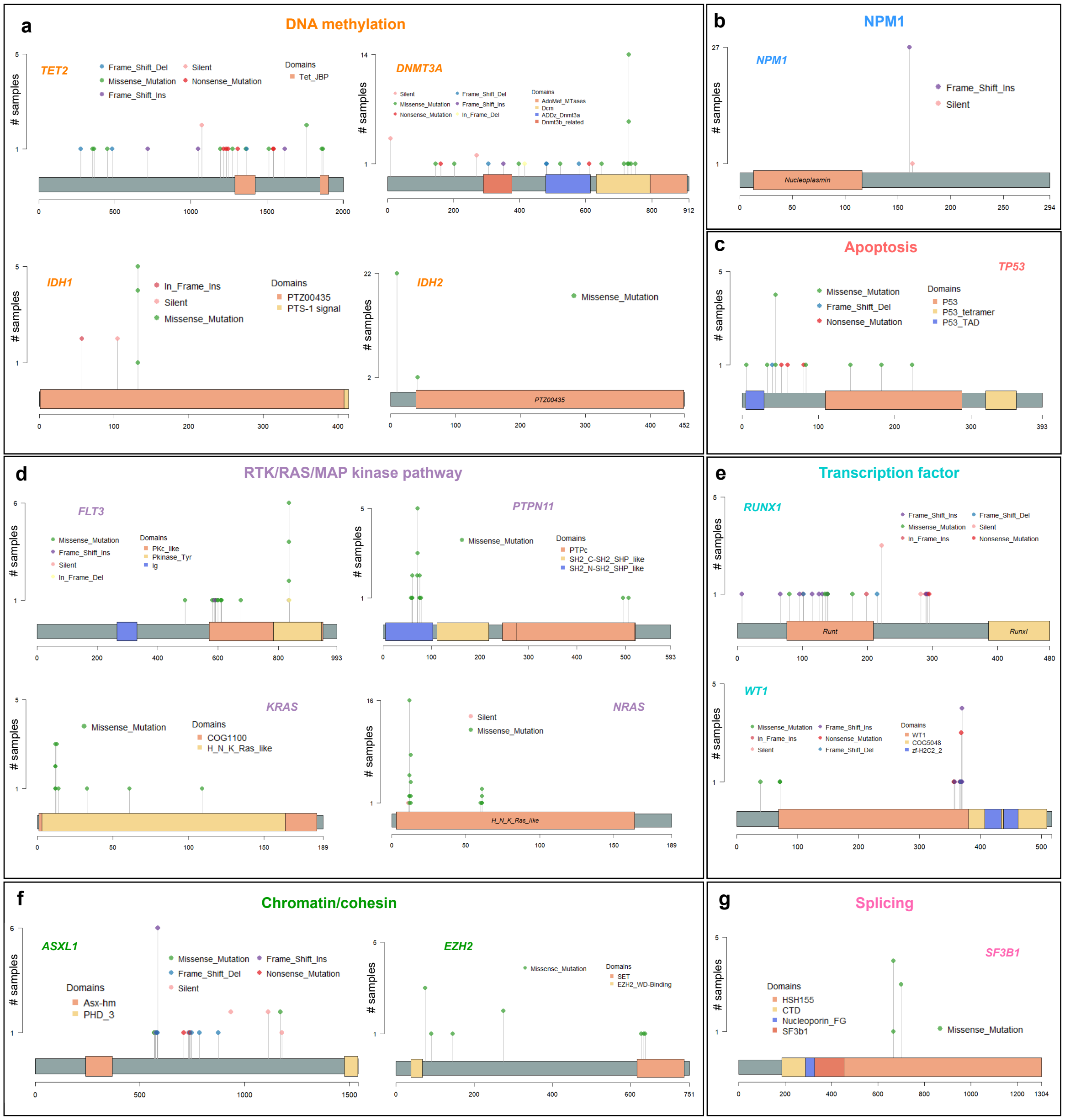


Supplementary Figure 6: Lollipop plots for variants detected by PHALCON in genes associated with different pathways: (a) DNA methylation (b) *NPM1* (c) apoptosis (d) RTK/RAS/MAP kinase pathway (e) transcription factor (f) chromatin/cohesin and (g) splicing.

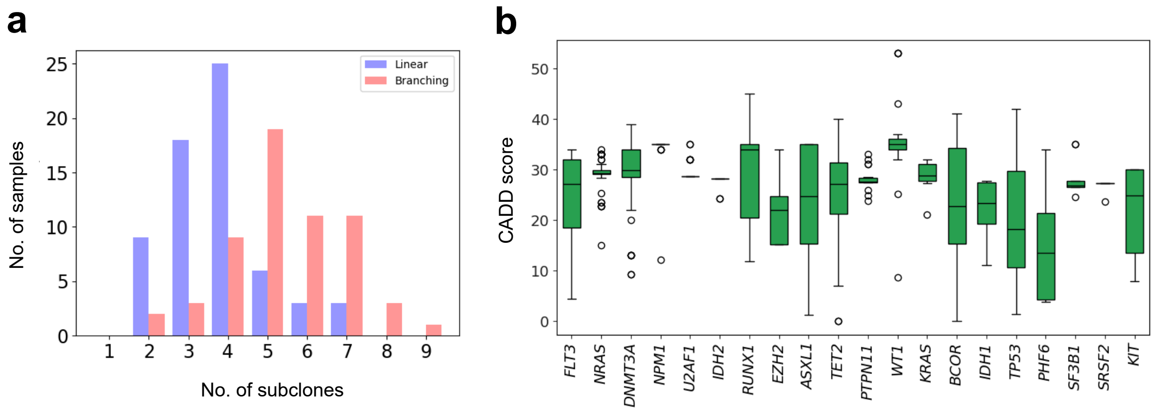


Supplementary Figure 7: (a) The distribution of the number of subclones inferred by PHALCON according to the evolution pattern (linear/branching) of the phylogenies reconstructed by PHALCON. (b) Boxplot of CADD scores of the mutations detected by PHALCON in the top mutated genes (top 20) in the AML cohort.

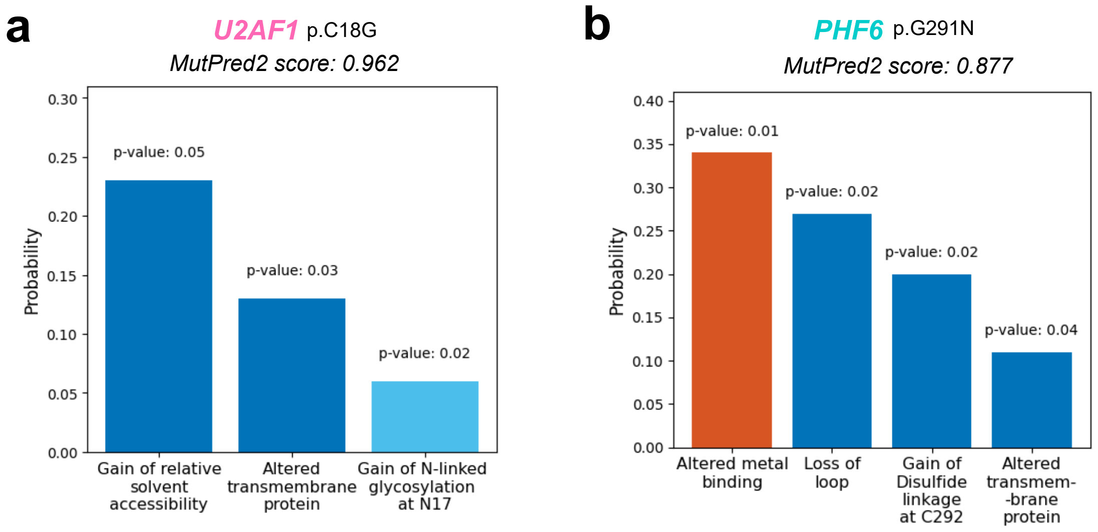


Supplementary Figure 8: MutPred2 prediction for significantly altered molecular mechanisms (p-value $< 0.05$) for (a) *U2AF1* mutation (p.C18G) and (b) *PHF6* mutation (p.G291N). The bars are colored based on the ontology of the molecular mechanism

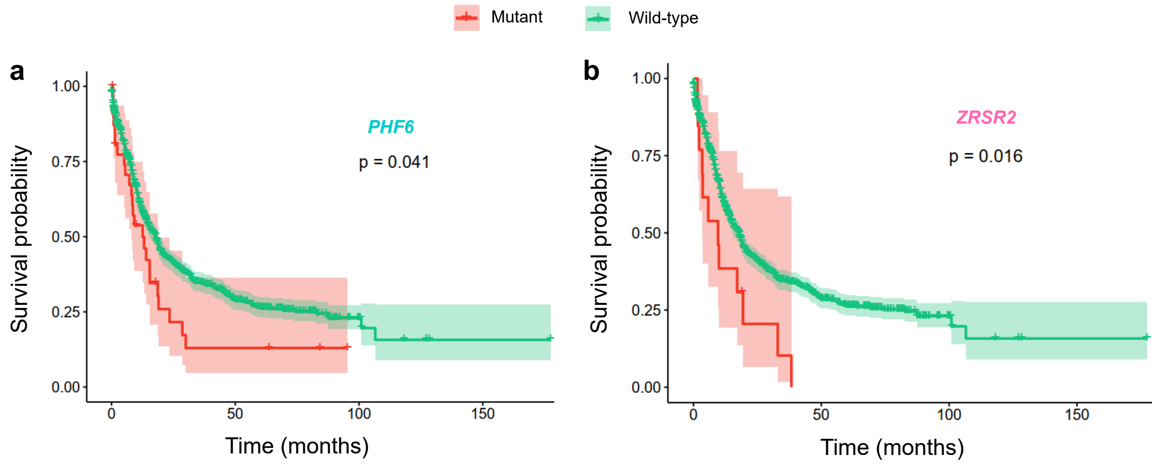


Supplementary Figure 9: Survival curves comparing mutant and wild-type groups for (a) *PHF6* and (b) *ZRSR2* in the combined TCGA AML and BeatAML 2.0 cohort.

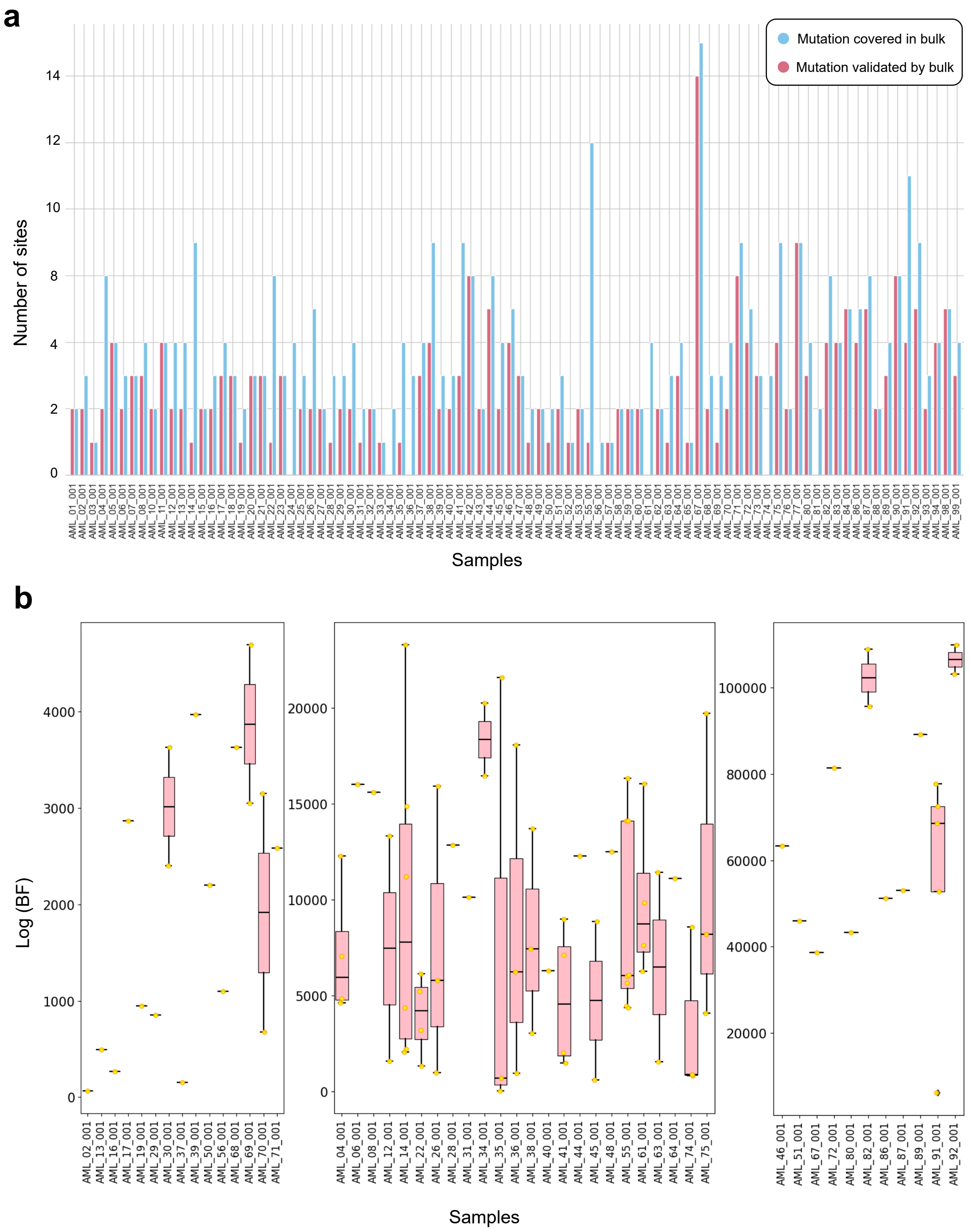


Supplementary Figure 10: (a) Barplot showing the number of PHALCON-detected variants orthogonally validated by bulk data. (b) Boxplot showing the log(Bayes Factor) values for mutations not having evidence in bulk data for each of the samples for which bulk data was available.

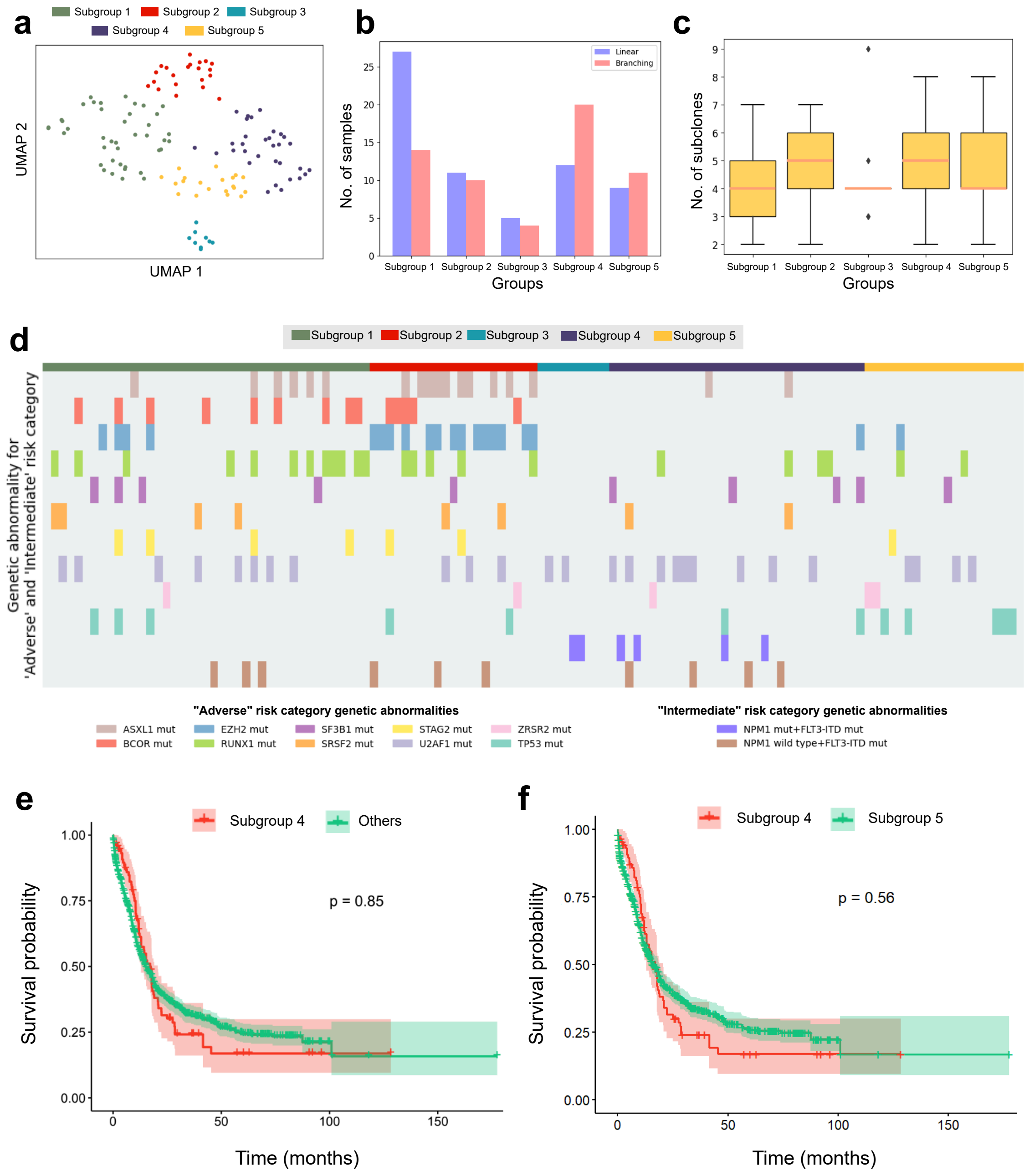


Supplementary Figure 11: (a) UMAP visualization of the pathway mutation profile of the five patient clusters. (b) Distribution of evolution pattern (linear/branching) for 5 subgroups in the AML cohort (123 patients) inferred based on the pathway mutation profiles of the samples derived by PHALCON. (c) Boxplot showing the number of subclones for samples belonging to each of the 5 subgroups. (d) Heatmap showing the presence of various genetic abnormalities across the subgroups for "adverse" and "intermediate" risk categories in AML based on ELN2022 [1] classification. The color bar on top indicates the subgroup association of each sample. (e) Survival curve comparing subgroup 4 and other subgroups in the combined TCGA AML and BeatAML 2.0 cohort. (f) Survival curve comparing subgroup 4 and subgroup 5 in the combined TCGA AML and BeatAML 2.0 cohort.

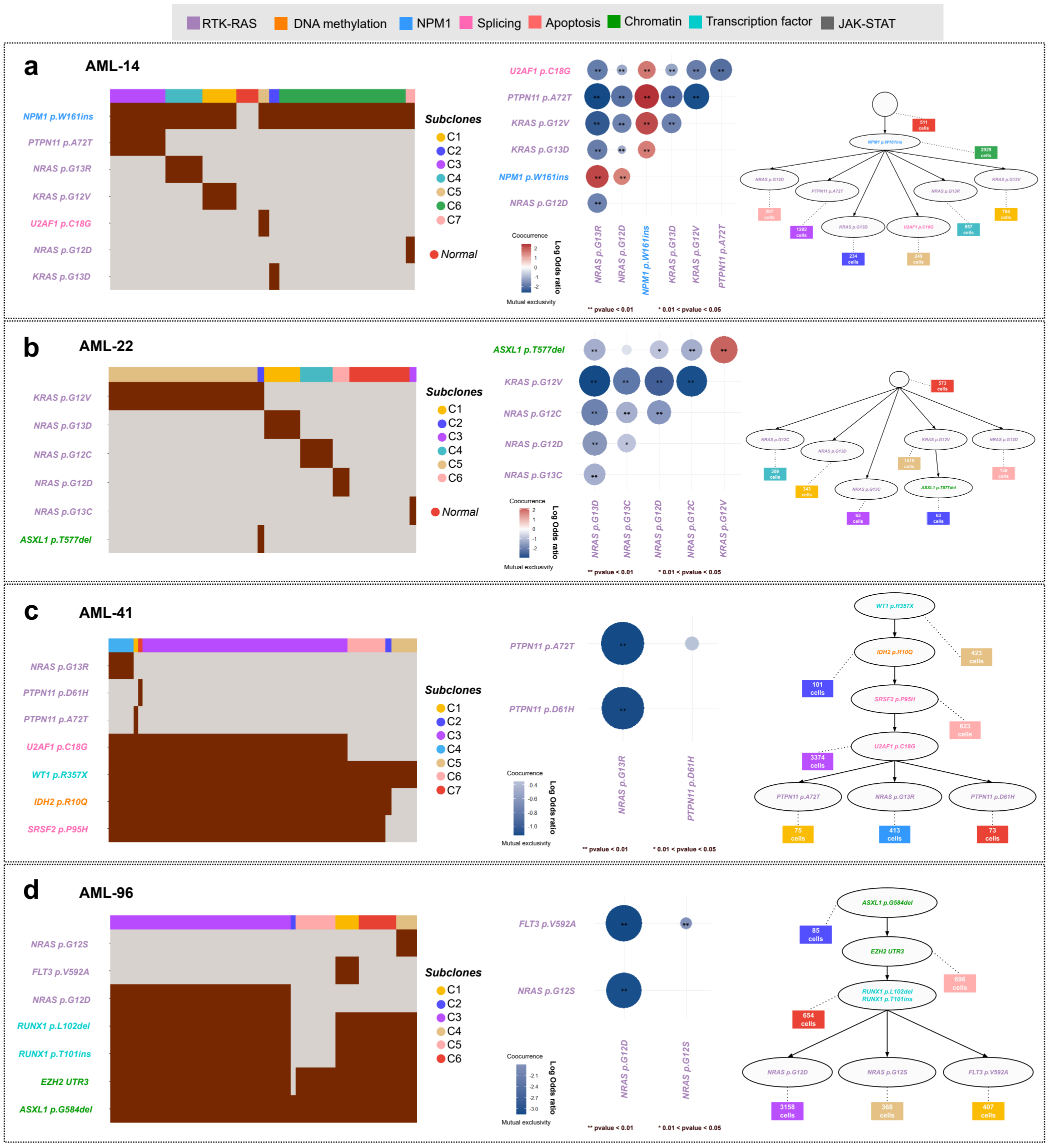


Supplementary Figure 12: Representative cases depicting cellular-level mutual exclusivity patterns of mutations in genes associated with the RTK/RAS/MAP kinase pathway for (a) AML-14 (b) AML-22 (c) AML-41 (d) AML-96. (left) Heatmap of clonally clustered mutations, illustrating the distribution of the driver mutations across different tumor clones. Each row represents a mutation and each column represents a cell. The clonal membership of individual cells is represented in the topmost row. (mid) Log odds ratio plot depicting the pairwise association between mutations in the patient sample. The color and size of a panel denote the magnitude of the logarithmic odds ratio (log OR). The reference bar maps the colors to their corresponding log OR values. Red denotes co-occurrence, while blue indicates mutual exclusivity. The statistical significance of these associations, based on p-values is marked by asterisks: * 0.01 < pvalue < 0.05, ** pvalue < 0.01. (right) Clonal phylogeny reconstructed by PHALCON showcasing the evolutionary history of each sample. The leaves (rectangular boxes colored according to clone identity) represent different sub-clones. Intermediate nodes contain sub-clonal mutations that are passed down the phylogeny. The root node contains the clonal mutations present across all cells. Genes are colored according to their associated pathways (grey box on top) across all three panels.

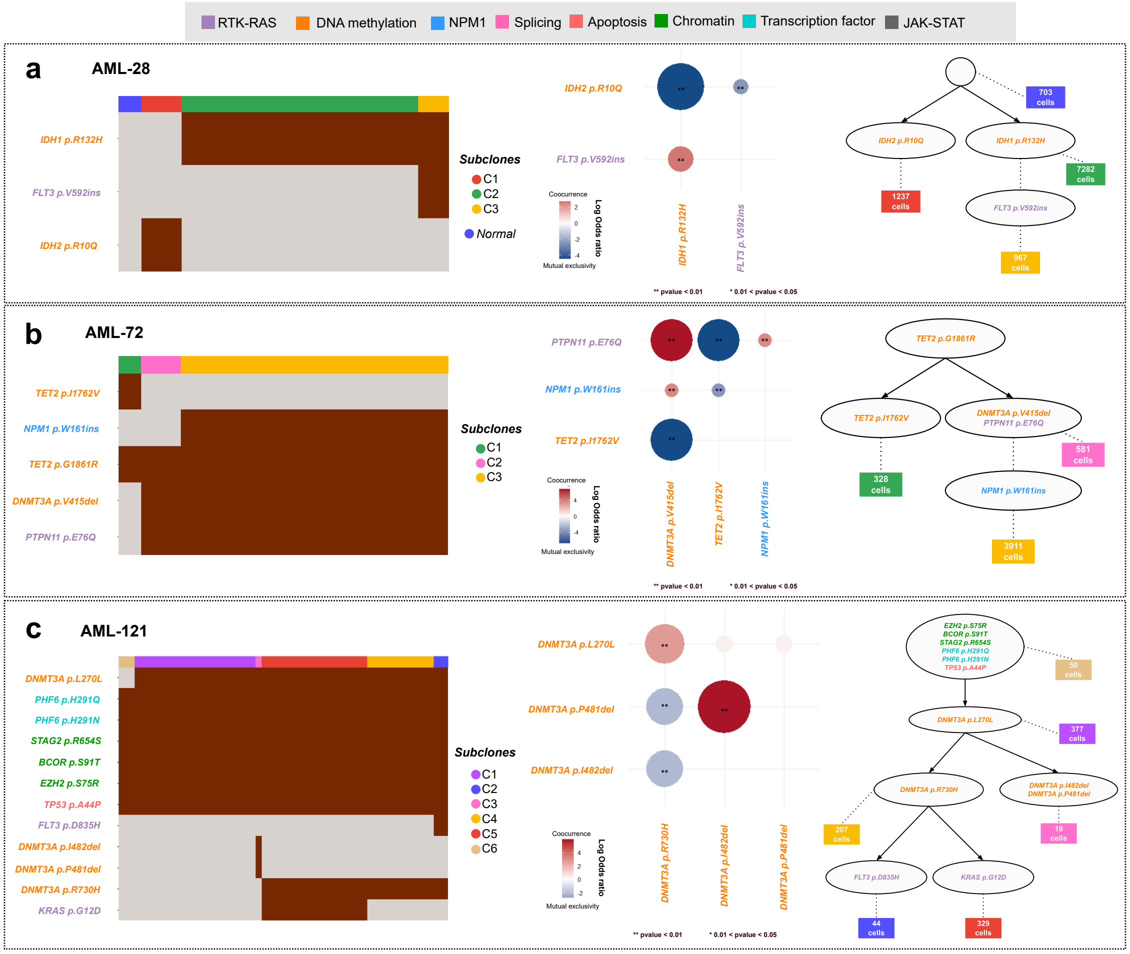


Supplementary Figure 13: Representative cases depicting cellular-level mutual exclusivity patterns of mutations in genes associated with the DNA methylation pathway for (a) AML-28 (b) AML-72, and (c) AML-121. (left) Heatmap of clonally clustered mutations, illustrating the distribution of the driver mutations across different tumor clones. Each row represents a mutation and each column represents a cell. The clonal membership of individual cells is represented in the topmost row. (mid) Log odds ratio plot depicting the pairwise association between mutations in the patient sample. The color and size of a panel denote the magnitude of the logarithmic odds ratio (log OR). The reference bar maps the colors to their corresponding log OR values. Red denotes co-occurrence, while blue indicates mutual exclusivity. The statistical significance of these associations, based on p-values is marked by asterisks: * $0.01 < \text{pvalue} < 0.05$, ** $\text{pvalue} < 0.01$. (right) Clonal phylogeny reconstructed by PHALCON showcasing the evolutionary history of each sample. The leaves (rectangular boxes colored according to clone identity) represent different sub-clones. Intermediate nodes contain sub-clonal mutations that are passed down the phylogeny. The root node contains the clonal mutations present across all cells. Genes are colored according to their associated pathways (grey box on top) across all three panels.

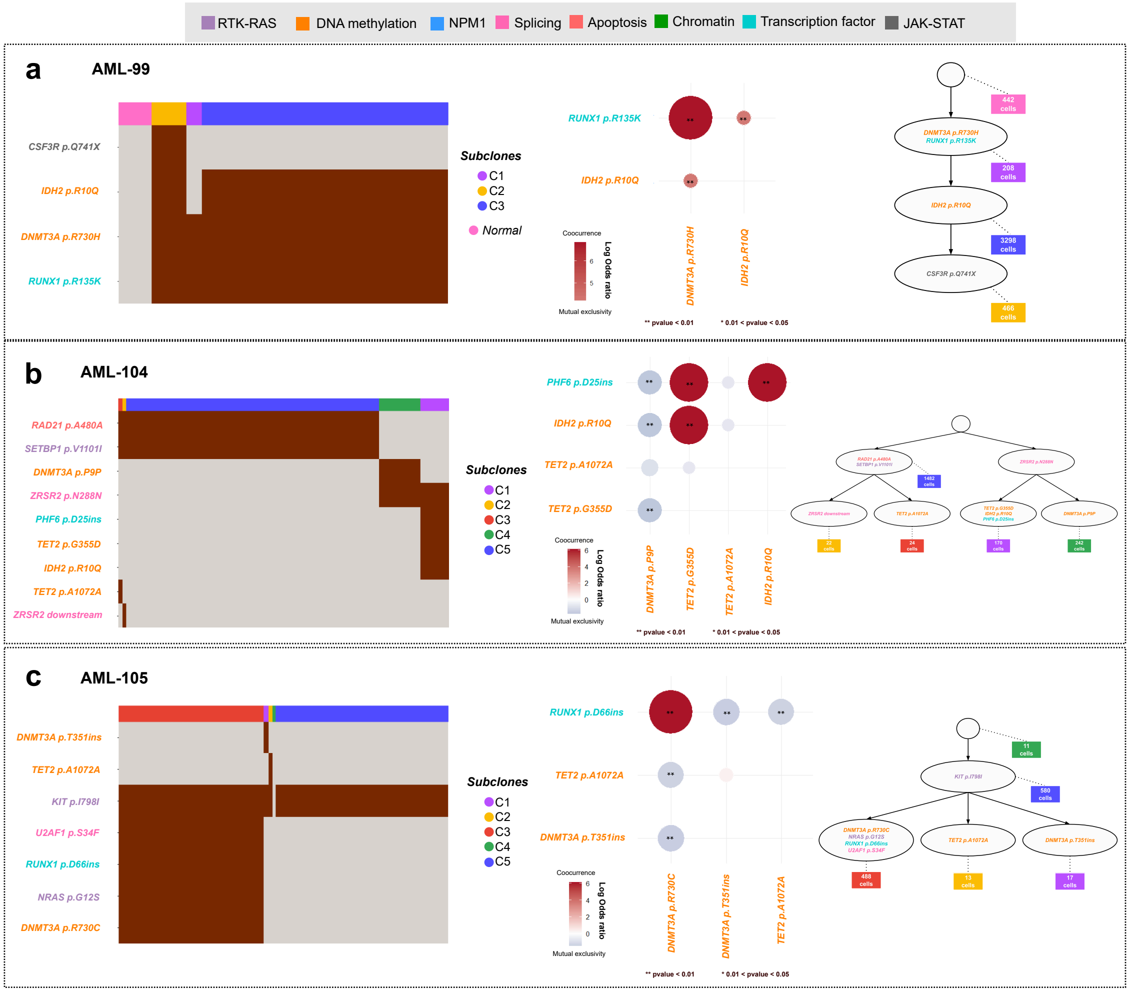


Supplementary Figure 14: Representative cases depicting cellular-level pairwise co-occurrence between genes associated with the DNA methylation pathway and Transcription factor pathway for (a) AML-99 (b) AML-104 (c) AML-105. (left) Heatmap of clonally clustered mutations, illustrating the distribution of the driver mutations across different tumor clones. Each row represents a mutation and each column represents a cell. The clonal membership of individual cells is represented in the topmost row. (mid) Log odds ratio plot depicting the pairwise association between mutations in the patient sample. The color and size of a panel denote the magnitude of the logarithmic odds ratio (log OR). The reference bar maps the colors to their corresponding log OR values. Red denotes co-occurrence, while blue indicates mutual exclusivity. The statistical significance of these associations, based on p-values is marked by asterisks: * $0.01 < \text{pvalue} < 0.05$, ** $\text{pvalue} < 0.01$. (right) Clonal phylogeny reconstructed by PHALCON showcasing the evolutionary history of each sample. The leaves (rectangular boxes colored according to clone identity) represent different subclones. Intermediate nodes contain sub-clonal mutations that are passed down the phylogeny. The root node contains the clonal mutations present across all cells. Genes are colored according to their associated pathways (grey box on top) across all three panels.

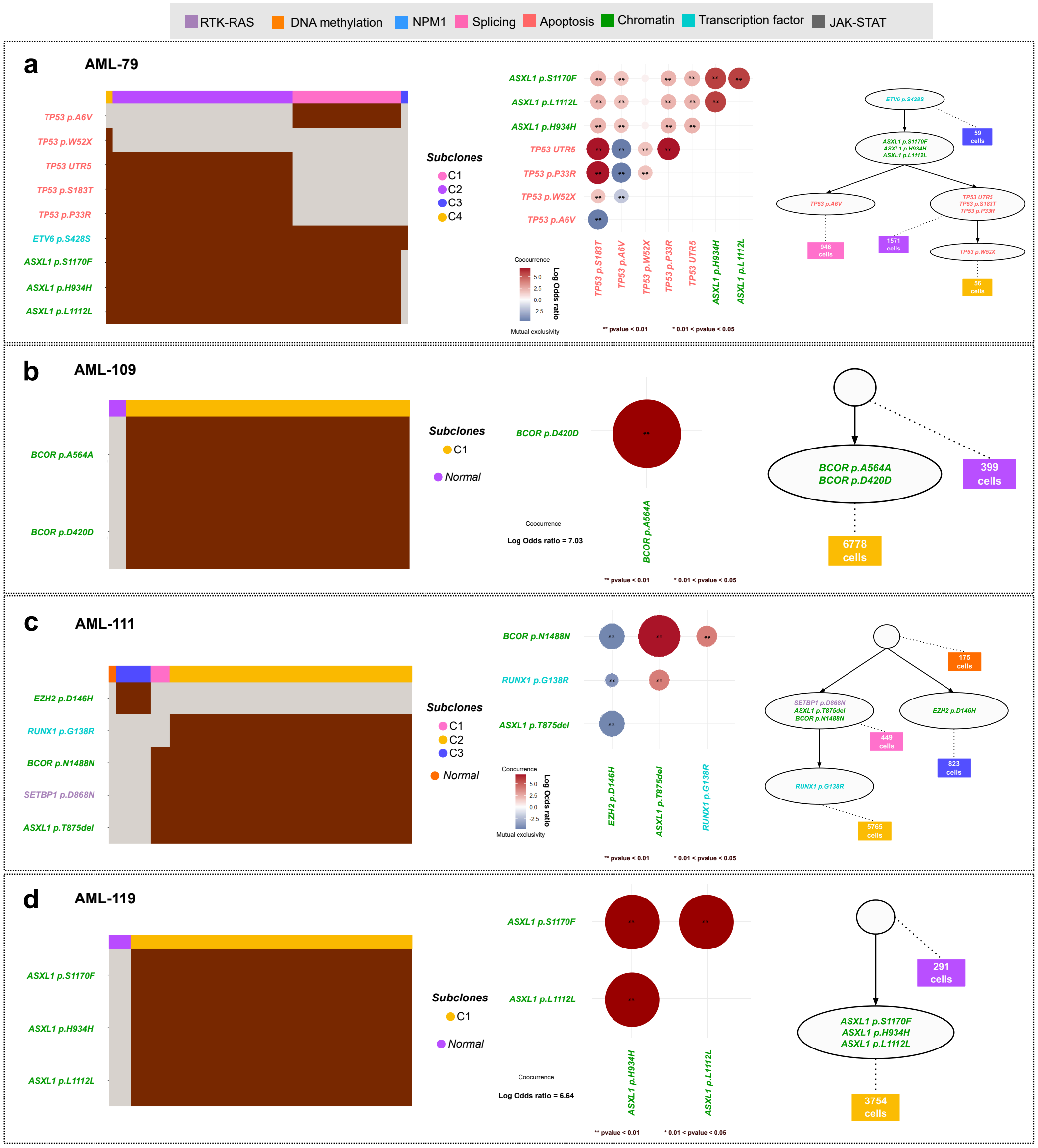


Supplementary Figure 15: Representative cases depicting cellular-level pairwise co-occurrence between genes associated with the Chromatin/cohesin pathway for (a) AML-79 (b) AML-109 (c) AML-111 and (d) AML-119. (left) Heatmap of clonally clustered mutations, illustrating the distribution of the driver mutations across different tumor clones. Each row represents a mutation and each column represents a cell. The clonal membership of individual cells is represented in the topmost row. (mid) Log odds ratio plot depicting the pairwise association between mutations in the patient sample. The color and size of a panel denote the magnitude of the logarithmic odds ratio (log OR). The reference bar maps the colors to their corresponding log OR values. Red denotes co-occurrence, while blue indicates mutual exclusivity. The statistical significance of these associations, based on p-values is marked by asterisks: * $0.01 < \text{pvalue} < 0.05$, ** $\text{pvalue} < 0.01$. (right) Clonal phylogeny reconstructed by PHALCON showcasing the evolutionary history of each sample. The leaves (rectangular boxes colored according to clone identity) represent different sub-clones. Intermediate nodes contain sub-clonal mutations that are passed down the phylogeny. The root node contains the clonal mutations present across all cells. Genes are colored according to their associated pathways (grey box on top) across all three panels.

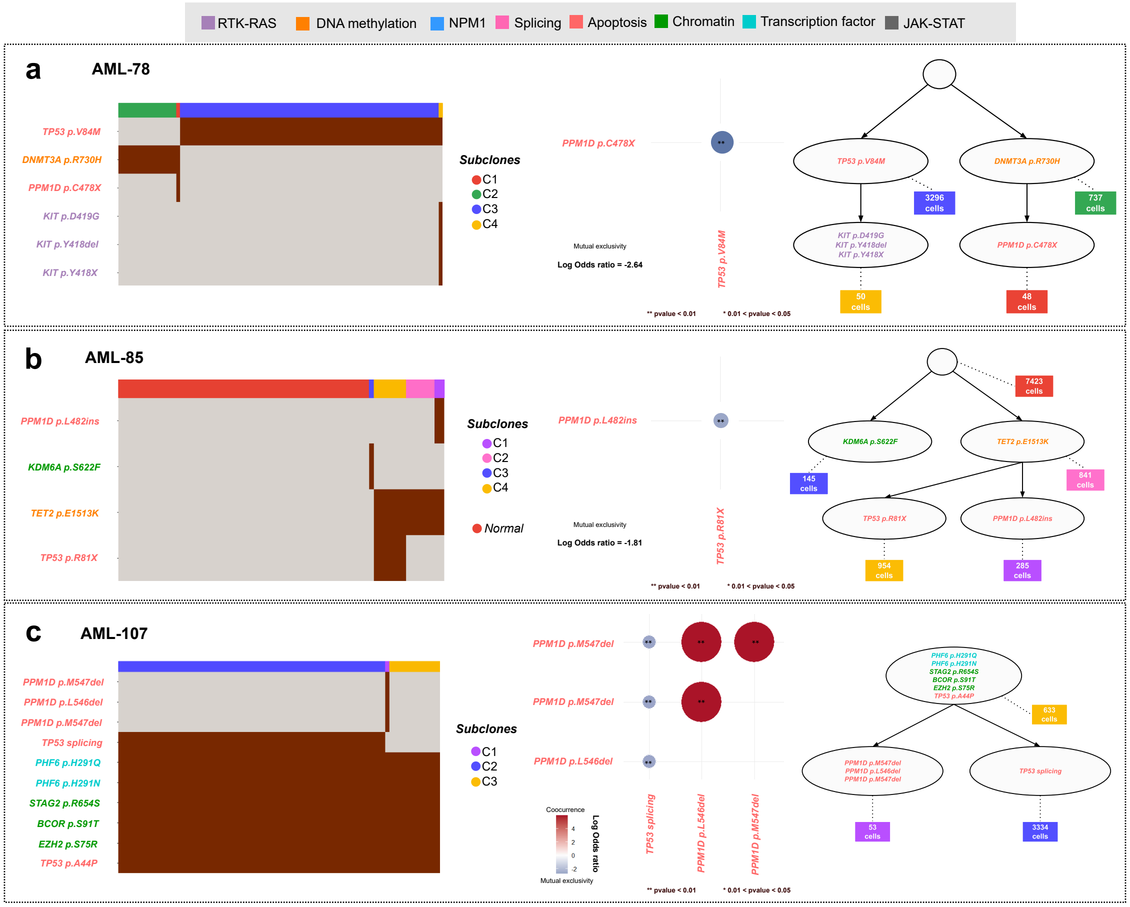


Supplementary Figure 16: Representative cases depicting cellular-level pairwise mutual exclusivity between genes associated with the Apoptosis pathway for (a) AML-78 (b) AML-85 and (c) AML-107. (left) Heatmap of clonally clustered mutations, illustrating the distribution of the driver mutations across different tumor clones. Each row represents a mutation and each column represents a cell. The clonal membership of individual cells is represented in the topmost row. (mid) Log odds ratio plot depicting the pairwise association between mutations in the patient sample. The color and size of a panel denote the magnitude of the logarithmic odds ratio (log OR). The reference bar maps the colors to their corresponding log OR values. Red denotes co-occurrence, while blue indicates mutual exclusivity. The statistical significance of these associations, based on p-values is marked by asterisks: * 0.01 < pvalue < 0.05, ** pvalue < 0.01. (right) Clonal phylogeny reconstructed by PHALCON showcasing the evolutionary history of each sample. The leaves (rectangular boxes colored according to clone identity) represent different sub-clones. Intermediate nodes contain sub-clonal mutations that are passed down the phylogeny. The root node contains the clonal mutations present across all cells. Genes are colored according to their associated pathways (grey box on top) across all three panels.

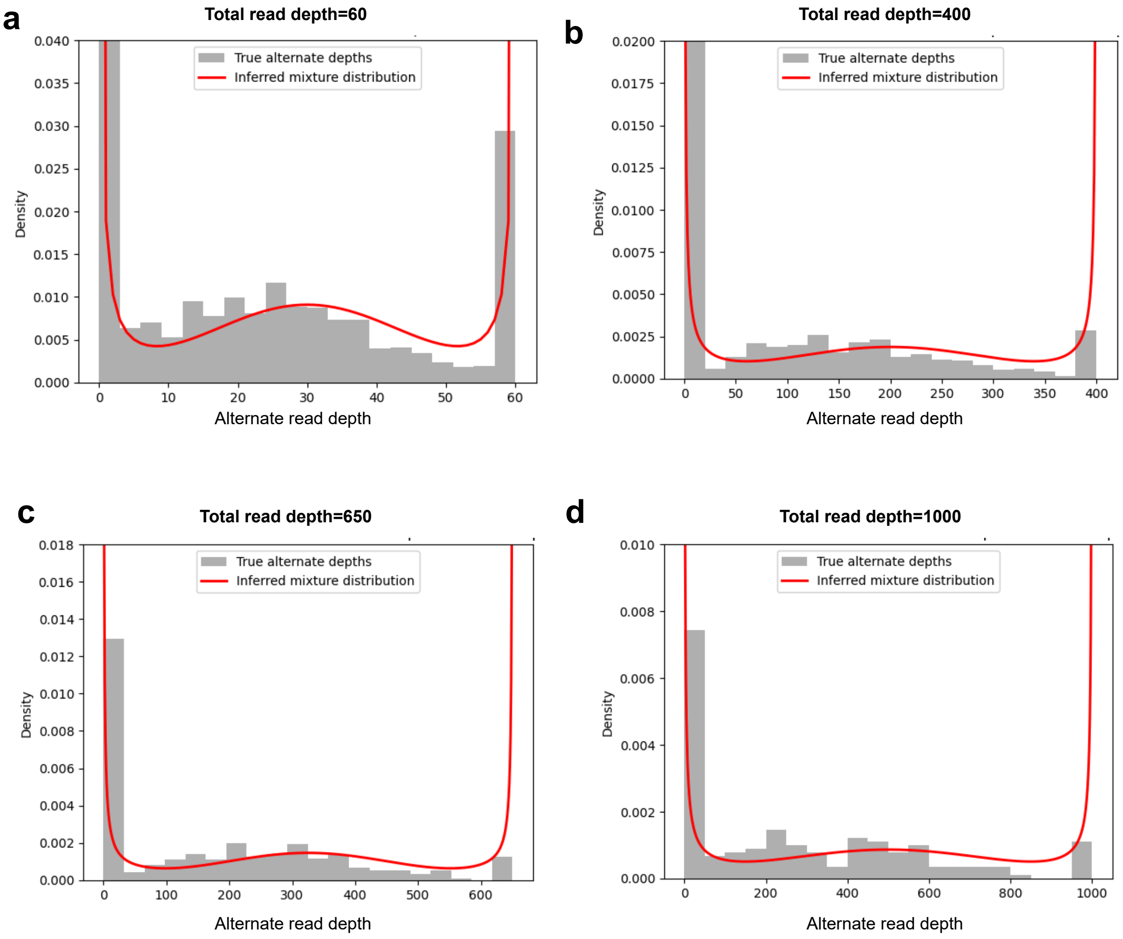


Supplementary Figure 17: Comparison of the distribution of true alternate depths and beta binomial mixture distribution used by PHALCON for confident heterozygous sites for varying total depths: (a) total read depth = 60, (b) total read depth = 400, (c) total read depth = 650, (d) total read depth = 1000. The sites were selected from the triple negative breast cancer datasets generated using MPT-seq [2]. Grey bars indicate the frequency of alternate depths for respective bins. The red curve indicates the probability density function inferred by PHALCON after using maximum likelihood estimation for shape, scale and weight paramaters. The y-axis has been accordingly adjusted as density.

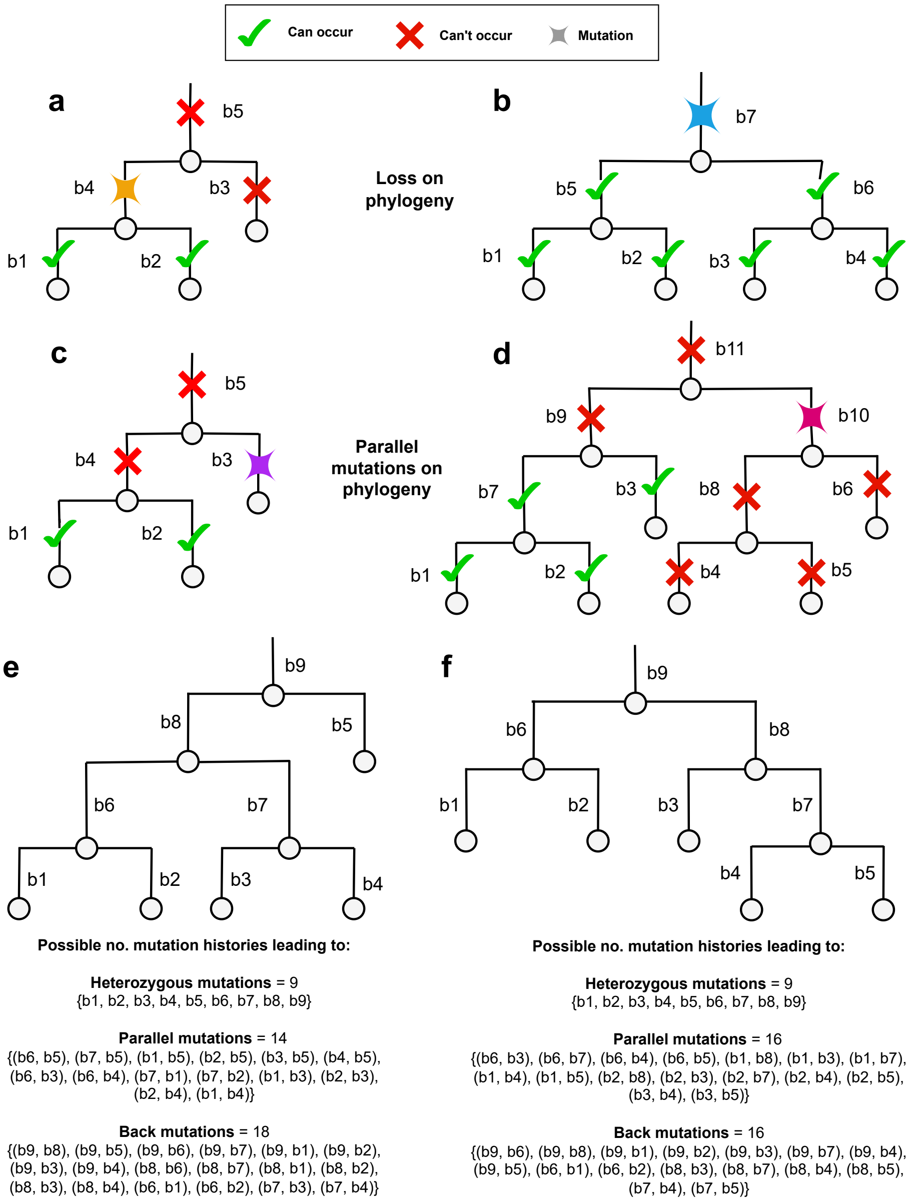


Supplementary Figure 18: Examples showing the possible mutation histories caused by heterozygous, parallel, and back mutations on the phylogeny. Branches are denoted using $b$ and numbered bottom-up (starting from the leaves, left to right). Nodes are represented using circles. (a)-(d) Examples of possible mutational history leading to mutation loss or parallel loss mutation. Both types of mutations involve a pair of branches. For mutational loss, the first instance denotes the branch where the mutation occurred, and the second instance denotes the loss of the same mutation. For parallel mutations, the first instance denotes the branch where a mutation occurred, and the second instance denotes the branch in an independent where the same mutation recurred. The first instance of the mutation is represented using a 4-pointed star. Green ticks denote branches where the second instance of the mutation is possible. A red cross on a branch denotes that the second instance can absolutely not occur at that branch. (e),(f) Examples of two different topologies with the same number of leaves, leading to different number of possible mutation histories leading to back mutations and parallel mutations.

### Supplementary Tables

Supplementary Table 1: Values of the tumor evolution parameters used for data simulation across different experimental settings

| **Experimental setting** | $p_{het}$ | $p_{back}$ | $p_{par}$ | $\gamma_{\textbf{clonal}}$ | $\gamma_{\textbf{clonal\_loss}}$ |
|---|---|---|---|---|---|
| Default | 0.9 | 0.05 | 0.05 | 0.3 | 0.05 |
| Increased back mutation | 0.8 | 0.15 | 0.05 | 0.3 | 0.1 |
| Increased parallel mutation | 0.8 | 0.05 | 0.15 | 0.3 | 0.05 |
| Increased back and parallel mutations | 0.7 | 0.15 | 0.15 | 0.3 | 0.05 |
| Infinite sites model | 1 | 0 | 0 | 0.3 | 0 |

Supplementary Table 2: Genes and biological pathways associated with the genes used for the derivation of pathway mutation profile

| **Gene name** | **Pathway** |
|---|---|
| *NPM1* | NPM1 |
| *DNMT3A* | DNA methylation |
| *IDH1* | DNA methylation |
| *IDH2* | DNA methylation |
| *TET2* | DNA methylation |
| *FLT3* | RTK/RAS/MAP kinase pathway |
| *KRAS* | RTK/RAS/MAP kinase pathway |
| *KIT* | RTK/RAS/MAP kinase pathway |
| *NRAS* | RTK/RAS/MAP kinase pathway |
| *PTPN11* | RTK/RAS/MAP kinase pathway |
| *PHF6* | Transcription factor |
| *RUNX1* | Transcription factor |
| *WT1* | Transcription factor |
| *ASXL1* | Chromatin/cohesin |
| *BCOR* | Chromatin/cohesin |
| *EZH2* | Chromatin/cohesin |
| *SF3B1* | Splicing |
| *SRSF2* | Splicing |
| *U2AF1* | Splicing |
| *TP53* | Apoptosis |

#### Supplementary Note 1: Description of filters used by PHALCON for identifying candidate mutation sites

##### Removal of Read Count Information in Cells based on Quality Score

This filter removes the read count data with an associated Genotype Quality score less than the given threshold. The Genotype Quality (GQ) score is derived from the Phred Likelihood(PL) value of the VCF file outputted by GATK variant caller. The GQ score indicates the difference in likelihood between the two most likely genotypes. A lower GQ value shows little confidence in picking one genotype over another. So, we use this filter to remove read count data where we cannot confidently infer one specific genotype. This filter can be turned off if the GQ scores are not available. Usually GQ scores are available with the loom file provided by Mission Bio Tapestri. By default, we use a threshold of 30 for this filter.

##### Removal of Read Count Information in Cells based on Read Depth

This filter removes the read count data from all the cells with a total read depth value less than the given threshold. This filter aims to remove very low coverage read count information from the analysis. We use a default value of 5 as a read depth threshold.

##### Removal of Read Count Information in Cells based on Variant Allele Frequency

For a diploid cell without any copy number alteration, a heterozygous mutation's variant allele frequency (VAF) should be around 0.5 (ideal value = 0.5, which can vary due to sampling noise). A much lower VAF value at a putative mutation site in a cell could be potentially indicating false-positives introduced during the whole genome amplification process of SCS. So, the read count information in such a locus in that cell with VAFs less than a certain threshold, is considered to be of low quality and hence removed by this filter. In the latter stages of filtering, for the putative mutation site, if there are not enough cells with available read count data, then such a variant site is considered to be a false-positive and is filtered out. This filter considers cells to harbor a putative mutation, if their mutation likelihood values are greater than 0.5. We use a default value of 0.2 as the variant allele frequency threshold.

##### Removal of Variants based on Read Count Information across all Cells

After applying the cell level filters above, if for a genomic site, there is a lack of a sufficient number of cells with high-quality read count information, then we cannot confidently determine if the site harbors a mutation. This filter removes the variant if the proportion of cells containing read count information across all cells is less than a certain threshold. For example, a threshold of 0.5 retains only the variants for which information is available in at least 50% of cells and hence variants with read count information in fewer than 50% of cells are removed. The default threshold value used for this filter is 0.5.

##### Removal of Variants based on the Fraction of Cells Mutated

If, for a genomic site, the number of cells harboring a mutation is very small, it is likely to be a false positive and needs to be removed. This filter considers only the fraction of cells with a mutation likelihood of at least 0.5. If this fraction is less than a given threshold at a variant site, then the variant is discarded. We use a default value of 0.004 as the threshold value. This value can be further reduced to lower thresholds to detect rare subclones. A significant number of false-positive variants could be included by lowering the filter threshold.

#### Supplementary Note 2: Description of possible mutation histories due to heterozygous, parallel, and back mutations for a given phylogeny

For a given phylogeny, PHALCON assigns the best mutation history at a site by selecting a branch (heterozygous) or pair of branches (parallel and back) for mutation placement that maximizes the log-likelihood. Here, we provide a detailed description of the various possible mutation histories for the events associated with PHALCON's finite-sites model. For mutation loss, if the first mutation occurs at branch $b$, and then it gets lost in the phylogeny at branch $b^-$, then the branch $b^-$ should strictly be present in the sub-tree rooted at $b$. For example, in Supplementary Fig. 18a, the mutation occurring at branch $b4$ can only be lost at branch $b1$ or $b2$. It cannot be lost at any adjacent branch, such as $b3$, nor at the ancestor branch, $b5$. If the mutation happens at the root branch, e.g. the mutation at $b7$ in Supplementary Fig. 18b, it is allowed to be lost at any other branch in the remaining topology.

For a parallel mutation, if we select a branch $b$ for placing the first occurrence, then the branch $b'$ for the second occurrence should be selected such that $b'$ is in an entirely different lineage than that of $b$. For example, in Supplementary Fig. 18c, the mutation placed at branch $b3$ can only occur recurrently on branch $b1$ or $b2$. It cannot occur at the ancestor branch $b5$; it cannot occur at the adjacent branch $b4$, as such an event can be explained by placing the same mutation at $b5$. Similarly, for another topology in Supplementary Fig. 18d, if a mutation occurs at branch $b10$, it can occur parallelly at four branches: $b1$, $b2$, $b3$, and $b7$, which are present in an entirely independent lineage and are not placed adjacently. The second mutation cannot occur on $b4$, $b5$, $b6$, or $b8$ as these are descendants of $b10$. $b9$ and $b10$ are also excluded for being the adjacent and ancestor branch respectively.

For a given topology $\mathcal{T}$ with $K$ leaves, there are a total of $2K - 1$ possibilities for a mutational history leading to a heterozygous mutation (heterozygous mutation can occur at exactly one branch). In contrast, for parallel mutations and back mutations, the total number of possible mutational histories is a function of $K$ and is entirely dependent on the topology. For example, for two different topologies with the same number of leaves ($K = 5$) shown in Supplementary Figs.

18e-f, the number of possible mutation histories leading to parallel, and back mutations vary depending on the topology, whereas the number of possible mutation histories leading to heterozygous mutation is $9 (= 2K - 1)$ for both the topologies.

#### Supplementary Note 3: Description of read count simulation

For different experimental settings, we used the values of $p_{het}$, $p_{par}$, $p_{back}$, $\gamma_{\text{clonal}}$ and $\gamma_{\text{non\_clonal}}$ as mentioned in Supplementary Table 1.

##### Simulation of Read Count data

After introducing dropouts, we generate the read counts at each genomic locus for each cell using the Pòlya urn model as described in [3] with some differences as detailed below.

We first simulate an artificial reference chromosome consisting of 40000 base pairs (bp). We then divide this chromosome into shorter chunks of 1000 bp for each of the individual cells. We then empirically fit a discrete distribution to the coverage values extracted from one of MissionBio's experimental datasets and randomly sample coverage values from this empirical probability distribution. To mimic the non-uniform coverage distribution of the SCS datasets, we further randomize the coverage values of individual genomic loci using a discretized Gaussian distribution.

To mimic the amplification process, we use the Pòlya urn model. For heterozygous positions, we start with a mutant allele and a wild-type allele in the urn. Whereas for dropout positions, we start with either a reference allele or an alternate allele. We randomly choose an allele from the urn, copy it, and return both the allele and its copy to the urn. With a probability of 0.25%, we select a genomic site to introduce False Positive (FP) errors, occurring due to erroneous amplification. At these chosen sites, we select a Polya urn round and return a different allele to the urn mimicking the faulty amplification. This Polya urn round is sampled from a linear probability mass function with a negative slope in the domain $[1, coverage + 1]$ where we set $P(coverage + 1) = 0$. This distribution is chosen to ensure that we have significant FP alleles introduced in the read counts at that location rather than a simple uniform distribution over $[1, coverage]$. This way of introducing FP errors makes the simulated data more similar to the real-time high-throughput SCS data. This process is repeated until we have simulated read counts in the urn equal to the coverage value sampled for that specific cell and site.

After the read counts are generated, sequencing errors are introduced. For introducing copy numbers, we randomly assigned copy numbers to the chunks of the artificial genome from the aneuploidy copy number population $\{1, 3, 4, 5, 6\}$ with respective probabilities $\{0.0625, 0.5, 0.25, 0.125, 0.0625\}$. The probability of choosing an additional copy starting from diploidy is lowered progressively, and the remaining probability is assigned to the loss of a copy. The probability of reference allele dropout is kept more ($cpn/(cpn-1)$, where $cpn$ is the copy number) than the probability of an alternate allele being dropped out. We synthesized two variations with copy number rates of 0.25 and 0.5. It is to note that the copy number rate here specifies the fraction of ground truth mutated sites, e.g., a copy number rate of 0.5 means that 50% of true mutated sites are affected by copy number (increase or decrease) and not 50% of the whole genome. The simulated dataset is converted into a read count matrix to use as input for PHALCON.

#### Supplementary References


[1] Döhner, H., Wei, A. H., Appelbaum, F. R., Craddock, C., DiNardo, C. D., Dombret, H., Ebert, B. L., Fenaux, P., Godley, L. A., Hasserjian, R. P., Larson, R. A., Levine, R. L., Miyazaki, Y., Niederwieser, D., Ossenkoppele, G., Röllig, C., Sierra, J., Stein, E. M., Tallman, M. S., Tien, H.-F., Wang, J., Wierzbowska, A., and Löwenberg, B. (2022). Diagnosis and management of aml in adults: 2022 recommendations from an international expert panel on behalf of the eln. *Blood*, 140(12):1345–1377.

[2] Leighton, J., Hu, M., Sei, E., Meric-Bernstam, F., and Navin, N. E. (2023). Reconstructing mutational lineages in breast cancer by multi-patient-targeted single-cell DNA sequencing. *Cell Genomics*, 3(1):100215.

[3] Singer, J., Kuipers, J., Jahn, K., and Beerenwinkel, N. (2018). Single-cell mutation identification via phylogenetic inference. *Nature Communications*, 9(1):5144.
